# The nucleus accumbens dorsomedial column: distinct anterior-posterior inputs and activation inversely associated with opioid relapse

**DOI:** 10.1101/2025.01.18.633747

**Authors:** Kenichiro Negishi, Vanessa I. Navarro, Mona S. Pishgar, Federica Ricciardi, Laura P. Montes, Lidice Soto Arzate, Josdell M. Guerra Ruiz, Diana Sotelo, Alejandro R. Toccoli, Huiling Wang, Yavin Shaham, Arshad M. Khan

## Abstract

Anatomical studies suggest that a dorsomedial region of nucleus accumbens (dmACB) forms a basal ganglia-like closed-loop circuit with medial ventral pallidum (mVP), lateral hypothalamic area (LHA), and paraventricular thalamic nucleus (PVT). The circuit’s topographic organization is unknown. We first used anterograde (*Phaseolus vulgaris* leucoagglutinin, PHAL) and retrograde (cholera-toxin B, CTB) tracers to map this organization. We then used immunoreactivity for Fos phosphorylated at serine 32 (pSer^32^-Fos-ir) as a proxy for the activation marker Fos (thus, we refer to pSer^32^-Fos hereafter as “Fos”) to determine whether the time-dependent increase (incubation) of oxycodone seeking after electric barrier-induced abstinence is associated with dmACB activation.

In Experiment 1, we injected rats with PHAL into the anterior PVT (aPVT) and CTB into either the mVP or the anterior LHA (LHAa) to determine the distribution of their connections within the dmACB. We also injected CTB into anterior and posterior dmACB to identify afferent inputs. In Experiment 2, we trained rats to self-administer oxycodone for two weeks and counted Fos-labeled cells in anterior and posterior dmACB, the surrounding ACB, and dorsal striatum regions after drug-seeking tests on either abstinence day 1 or day 15 following electric barrier-induced abstinence.

PHAL-labeled aPVT axons and CTB-labeled mVP and LHA neurons were confined to dmACB, forming a longitudinal column along the anterior-posterior axis of ACB. Inputs to anterior and posterior dmACB came from different cortical and subcortical regions. dmACB Fos+ cell counts were higher after oxycodone-seeking tests on day 1 than on day 15 and were negatively correlated with oxycodone seeking.

These results characterize a connection-defined dmACB circuit and show that incubation of opioid-seeking is unexpectedly associated with reduced dmACB activation.

## Introduction

Cerebral hemisphere connections integrate internal and external sensory information to guide motivated behaviors, which are thought to depend on basal ganglia circuits (Alexander et al., 1986). These circuits are traditionally described as parallel, largely non-overlapping closed-loop pathways spanning major forebrain divisions above the hindbrain (Foster et al., 2021; Lee et al., 2020). However, most classical basal ganglia circuit models exclude the hypothalamus, despite its central role in motivated behaviors (Hoebel & Teitelbaum, 1962; Margules & Olds, 1962; Stuber & Wise, 2016). A broader synthesis incorporates key hypothalamic structures into basal ganglia frameworks (Swanson, 2000). This notion guided the identification of amygdala-and frontal cortex-centered basal-ganglia circuits interacting with specific hypothalamic regions (Barbier et al., 2020; Hintiryan et al., 2025).

Evidence suggests the existence of a basal ganglia-like circuit involving regions connected to the *lateral hypothalamic area (Nissl, 1913)* (LHA). A study using anterograde- and retrograde-tracer injections identified a closed-loop circuit comprising the *infralimbic area (Rose & Woolsey, 1948)* (ILA), dorsomedial *accumbens nucleus (Ziehen, 1897–1901)* (dmACB), medial ventral pallidum (mVP), anterior LHA (LHAa; also called a posterior part of lateral preoptic area), and anterior *paraventricular thalamic nucleus (>1840)* (Thompson & Swanson, 2010) (aPVT). However, the circuit’s anterior-posterior organization, a critical feature of basal ganglia circuitry, has not been investigated.

Recent findings identify the dmACB as a specialized striatal domain. Several studies report high expression of hybrid dopamine receptor subunit (D1R-D2R) co-expressing neurons (Gokce et al., 2016; He et al., 2021; Saunders et al., 2018; Stanley et al., 2020), dopamine D3R-expressing neurons (Gagnon et al., 2017; Gangarossa et al., 2013), and the striosomal neuropeptide marker, prepronociceptin (Hueske et al., 2024), in dmACB. These data indicate that the dmACB is molecularly and anatomically distinct from neighboring striatal regions, suggesting that it is part of a discrete basal ganglia-like circuit.

Basal ganglia circuits are difficult to define anatomically due to their complex topographic organization. Top-down approaches (tracer injections into cortical or striatal areas) were instrumental in mapping corticostriatal and striatopallidal projections (Foster et al., 2021; Hintiryan et al., 2016; Lee et al., 2020). However, the widespread distribution of corticostriatal axons can obscure functionally distinct striatal domains (Berendse et al., 1992; Vertes, 2004). In contrast, bottom-up methods leverage the relative simplicity and focal nature of striatopallidal and striatohypothalamic projections, which can identify discrete connectivity motifs (Heimer et al., 1991; Li et al., 2025; Zahm & Heimer, 1993). Here, we used both top-down and bottom-up anatomical strategies to further characterize a basal ganglia-like circuit along the anterior-posterior axis of dmACB (first described in the previous version of this preprint using pilot experiments). We also determined whether neuronal activity of dmACB is associated with relapse to opioid seeking, which extends the studies presented in the previous version of this preprint with new behavioral data.

In Experiment 1, we used anterograde (PHAL) (Gerfen & Sawchenko, 1984) and retrograde (CTB) (Luppi et al., 1990) tracers to define the connectivity of dmACB. We first injected tracers into aPVT (PHAL), mVP (CTB), and LHAa (CTB) to determine the spatial extent of the dmACB. We then used retrograde tracing (CTB injections in dmACB) to determine whether anterior versus posterior dmACB are part of distinct forebrain and midbrain projection networks.

In Experiment 2, we assessed the behavioral relevance of dmACB activation using the neuronal activation marker Fos. We used a rat model of time-dependent increases (“incubation”) of relapse to opioid seeking following electric barrier-induced abstinence (Fredriksson et al., 2020). In this model, abstinence is induced by increasing electric-barrier intensity until rats cease drug self-administration; relapse is then evaluated without the barrier and the drug (Fredriksson et al., 2021b; Negishi et al., 2024). Using oxycodone as the self-administered opioid, recent work has implicated the *claustrum (Burdach, 1822)* (CLA), *orbital region (Rolando, 1819)* (i.e., orbitofrontal cortex, ORB), dorsomedial striatum, and *subiculum ventral part (Swanson & Cowan, 1975)* (SUBv) in this form of incubation (Fredriksson et al., 2021a; Fredriksson et al., 2023; Ma et al., 2024; Negishi et al., 2025). Here, we tested whether neuronal activation in anterior versus posterior dmACB is associated with incubation of oxycodone seeking.

## Materials and Methods

### Experiment 1: Anatomical characterization of the dmACB column

#### Overview

In pilot studies, we used four male rats to begin anatomically characterizing the dmACB column, as described in the previous version of this preprint. We used methods like those in Experiment 1 and describe them in detail in that version (see the previous version of this preprint for histological and microscopy procedures used in **Figs. 9–12**). We designed Experiment 1 to confirm and extend these pilot results in a larger cohort and to ensure their reproducibility and update this newest preprint version.

#### Subjects

In Experiment 1, we used male (n=26) and female (n=49) Sprague-Dawley rats (Charles River) weighing 180–430 g. We maintained rats under a reverse 12:12 h light/dark cycle (lights off at 8:00 A.M.) with free access to food and water. We housed two rats per cage prior to surgery and individually after surgery. We performed Experiment in accordance with the NIH *Guide for the Care and Use of Laboratory Animals (8th edition)*, under protocols approved by the Animal Care and Use Committees of NIDA Intramural Research Program and the University of Texas at El Paso.

#### Intracranial surgery

We used the iontophoresis method for PHAL and CTB co-injections. We anesthetized the rats and maintained them under isoflurane (5% induction, 2–3% maintenance, Covetrus). We head-fixed the rats on a stereotaxic instrument and made midline incisions to prepare the scalp prior to craniotomy. We exposed the surface of the dura mater with a round head drill and used a syringe tip to remove the dura at the brain entry site. We prepared tracer mixtures by dissolving 2.5% PHAL (Vector Laboratories; Cat# L-1110-5) and 0.25% CTB (List Labs; Cat# 104) in filtered 1× PBS. For iontophoresis, we loaded tracers into glass micropipettes and lowered them into brain targets based on coordinates from Paxinos and Watson (2014). We used a BAB-600 (M1160; Kation Scientific) to deliver 5 µA in 7 s ON/OFF cycles over 15 min. We removed the micropipettes after 10 min to allow diffusion. We used 4 groups of rats (n=10–25 per group) and injected the tracers’ mixture unilaterally into dmACB (n = 16; AP: +1.0–2.0 mm, ML: ±0.8 mm, DV: −6.2 mm from dura), mVP (n = 25; AP: 0.1 mm, ML: ±1.8 mm, DV: −7.2 mm from dura), LHAa (n = 24; AP: −1.5 mm, ML: ±2.0 mm, DV: −8.2 mm), and aPVT (n = 10; AP: −1.5 mm, ML: ±1.0 mm, DV: −5.1 mm from dura; angled 10° inward). We provided the rats with postsurgical care and euthanized them 7–10 days after surgery.

#### Tissue preparation

We deeply anesthetized rats in an enclosed glass desiccator saturated with isoflurane vapor for 70 s and perfused them transcardially at room temperature with 200 mL of PBS, pH 7.4 at room temperature, followed by 350 mL 4% paraformaldehyde dissolved in PBS. We removed the brains and placed them into the PFA solution for post-fixation overnight at 4°C. We transferred the brains to PBS containing 30% sucrose on the following day. We froze the brains in dry ice and sectioned them using a cryostat (Leica Microsystems). We collected coronal sections of 40-µm thickness into four series and stored them in cryoprotectant at −20°C (50% 2× PBS, 20% glycerol, and 30% ethylene glycol) until further processing.

#### Immunohistochemistry (IHC) for PHAL and CTB

To single-label PHAL and CTB, we used immunofluorescence based on our previous work (Bono et al., 2025). We removed tissues from cryoprotectant and washed them in 1× PBS (pH 7.4 at room temperature) for three washes, each for 10 min (3 × 10). We incubated tissues for 2 h in a blocking solution containing 2% (v/v) normal donkey serum (EMD Millipore; Catalog #S30) and 0.3% Triton X-100 (Sigma-Aldrich; Catalog #T8532). We then incubated the tissues in a primary antibody cocktail containing antibodies raised against PHAL (host: rabbit; dilution: 1:4,000; Vector Labs, Cat # AS-2224; RRID: AB_2313686) and CTB (host: goat; dilution: 1:10,000; List Biological Cat # 703; RRID: AB_10013220) in blocking solution for 24 h in 4°C. After 3 × 10 PBS washes, we incubated the sections in a secondary antibody cocktail with an AlexaFluor 555-conjugated antibody raised against rabbit (host: donkey; dilution: 1:500; Jackson ImmunoResearch Labs, Cat# 711-565-152; RRID: AB_3095471), an AlexaFluor 488-conjugated antibody raised against goat (host: donkey; dilution: 1:500; Jackson ImmunoResearch Labs, Cat# 705-545-147; RRID: AB_2336933), and NeuroTrace 660/640 (dilution: 1:200; ThermoFisher Scientific, Cat# N21483) for 5 h. After a final wash in PBS, we mounted the free-floating sections onto glass slides and left them to dry at room temperature for 2 h. We coverslipped tissues using slightly diluted DAPI-Fluoromount-G (Electron Microscopy Sciences, Cat# 17984-24).

#### Antibody validation

The primary antibody cocktail solution for this study only contained antibodies raised against PHAL and CTB. We did not observe specific staining for PHAL or CTB in rats that did not receive tracer injections. Additionally, we did not observe specific staining for injection cases that did not effectively transport tracers (n=12). The same validation approach and outcome were found in other reports that used the same antibodies (Hahn & Swanson, 2010, 2012). We used no-primary controls to determine the specificity for secondary antibodies. We performed the IHC procedures as described above but with the omission of primary antibodies in the first cocktail. We did not detect any labeling for the secondary antibodies we used in the absence of the primary antibodies.

#### Microscopy and image processing

For immunofluorescence imaging, we used an Olympus VS200 slide scanner. We obtained four-channel mosaic images for each section using a ×10 objective lens and then exported images for the channels separately as grayscale 16-bit PNG-formatted files. We then used a custom Python script to apply linear scaling to produce 8-bit images with upper and lower bounds based on their intensity histograms. We obtained pseudo-Nissl stains by inverting the Neurotrace images.

#### Segmentation and mapping of immunohistochemically detected tracers

The atlas-based mapping approach we used follows the cytoarchitectonic approach described by Swanson (1992, 1998, 2004, 2018). Briefly, we obtained formal cytoarchitecture-based definitions and nomenclature for gray matter regions for all areas examined (see Table C in Supplementary Information Folder 1 from Swanson (2018)). We drew boundaries based on Nissl-stained reference sections that we superimposed on images of labeled tracers for localization. We performed Nissl- and immuno-staining on the same sections, which allowed direct image alignment as separate layers using Adobe Illustrator.

To obtain unbiased and reliable image segmentation, we used a custom Python script to generate vector segmentations for CTB and PHAL labeling. We trained a neural network to predict CTB-positive neurons using Cellpose-SAM (Pachitariu et al., 2025; Stringer & Pachitariu, 2025), and converted outputs to SVG format to facilitate alignment on Adobe Illustrator. For PHAL axon segmentation, we trained a basic U-Net model (Ronneberger et al., 2015) to predict labeled axons. We then converted the pixel-wise predictions into 8-bit grayscale images and applied a threshold to obtain binarized bitmap images. We used Adobe Photoshop to convert binary images into Illustrator-compatible work paths. Finally, we aligned segmented neurons and axons to their original CTB and PHAL images for proofreading. The vector format facilitated selection, deletion, and addition of individual objects using direct selection tools in Adobe Illustrator.

We then transferred the CTB and PHAL labeling to standard digital atlas plates from *Brain Maps 4.0* (BM4; Swanson, 2018) and performed manual atlas registration to obtain maps. For atlas registration, we first transferred segmentation objects together with boundary parcellations to their corresponding BM4 digital atlas plates. We applied affine transformations (linear scaling and rotations) to get a first approximation and then used the *Envelope Distort* tool to achieve the non-linear transformations required for precise registration. Matching histological sections to atlas levels involved careful observation of Nissl-based boundaries, histological plane-of-section (Zséli et al., 2016) and other forms of non-linear distortion that result from tissue processing (Simmons & Swanson, 2009). Accordingly, our maps show data from the best-matched tissue sections (and sometimes just portions of them), for each atlas plate.

#### Intersection maps

To identify spaces of convergent connectivity, we first generated kernel density estimates (KDE) from atlas-mapped CTB and PHAL data. We exported CTB objects as SVG files and rasterized and exported PHAL objects as PNG files. We used a custom Python script with the SciPy stats module (Virtanen et al., 2020) to apply the *gaussian_kde* class to estimate continuous probability density functions, which were then evaluated on a regular grid to obtain discretized density maps. We used the following function to generate intersection scores:

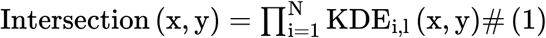

In this general form, the intersection map is defined as the product of density maps from specified brain connections. Here, KDE_i1_(x,y) denotes the discretized kernel density estimate (KDE) for label *l* from region *i*, evaluated at atlas coordinates *(x,y)*. The resulting maps were formed by element-wise multiplication of each label in atlas grid space. We adopted this approach to define and visualize dmACB and its corresponding mVP region using the following equations:

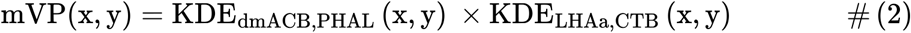

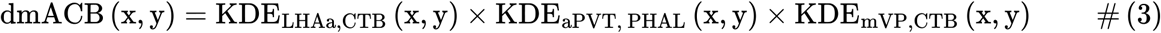

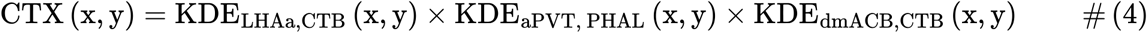

We treated the outputs from equations (2)–(4) as continuous intersection density maps and rescaled them for visualization and thresholding. We computed a global scaling factor by multiplying the sum of all grid values by the total grid area. To identify credible region boundaries, we sorted grid values into a one-dimensional array in descending order and computed their cumulative sum. We approximated a 95% threshold by identifying the threshold value at which the cumulative sum reached 95% of the total cumulative density. For visual clarity, we exported the intersection maps with a gamma-adjusted color scale (gamma=0.5) after computing the credible region thresholds. We then aligned the resulting intersection maps and credible region boundaries to their corresponding atlas plates.

#### Connection density quantification

We quantified connection density by generating regions of interest (ROIs) and using them as masks to measure kernel density estimate (KDE) values on a common grid. We manually outlined the striatum from *Brain Maps 4.0* atlas plates to obtain striatal ROIs. We defined dmACB ROIs using the 95% credible region approach described for the intersection maps. We converted both striatal and dmACB contours into grid-aligned binary masks using NumPy (Harris et al., 2020). Using the same grid, we computed KDE-based connection densities for each label. We restricted these densities to each ROI by multiplying them element-wise with the corresponding binary masks, then summed the values to obtain total connection densities for the striatum and dmACB. For each label, we calculated the proportion of striatal connection density contained within the dmACB as the ratio of dmACB density to total striatal density.

#### Anatomical nomenclature

For the anatomical data, we adopted the spatial framework, nomenclature, and ontology of Swanson (Swanson, 2015; Swanson, 2018) **(Table 1)**. We report brain regions using the naming conventions of Swanson (2015), as implemented for the rat brain in *Brain Maps 4.0* (Swanson, 2018), and we use standard terms throughout. We italicize each standard term and include the associated (author, date) citation that first defined it (e.g., *lateral hypothalamic area (Nissl, 1913)*). When priority could not be determined, the year “(>1840)” was assigned to indicate that the term was defined after 1840, approximately marking the introduction of cell theory (e.g., *dorsomedial hypothalamic nucleus (>1840)*).

**Table 1:** List of anatomical abbreviations.

|  |  |
|---|---|
| AAA | *anterior amygdalar area (Gurdjian, 1928)* |
| ACA | *anterior cingulate area (Brodmann, 1909)* |
| ACAd | *anterior cingulate area dorsal part (Krettek & Price, 1977)* |
| ACAv | *anterior cingulate area ventral part (Krettek & Price, 1977)* |
| ACB | *accumbens nucleus (Ziehen, 1897–1901)* |
| ACBcore | accumbens nucleus core |
| aco | *olfactory limb of anterior commissure (>1840)* |
| AD | *anterodorsal thalamic nucleus (>1840)* |
| a-dmACB | dorsomedial column of accumbens nucleus, anterior part |
| AHN | *anterior hypothalamic nucleus (>1840)* |
| AHNa | *anterior hypothalamic nucleus anterior part (>1840)* |
| AHNc | *anterior hypothalamic nucleus central part (>1840)* |
| AHNp | *anterior hypothalamic nucleus posterior part (>1840)* |
| AI | *agranular insular area (Krettek & Price, 1977)* |
| alv | *alveus (Burdach, 1822*) |
| AM | *anteromedial thalamic nucleus (>1840)* |
| amc | *amygdalar capsule (Swanson, 1998)* |
| aPVT | *anterior paraventricular thalamic nucleus (>1840)* |
| AUDd | *dorsal auditory areas (Sally & Kelly, 1988)* |
| AUDpo | *posterior auditory area (Doron et al., 2002)* |
| AV | *anteroventral thalamic nucleus (>1840)* |
| BLAa | *basolateral amygdalar nucleus anterior part (>1840)* |
| BLAp | *basolateral amygdalar nucleus posterior part (>1840)* |
| BLX | *basolateral amygdalar complex (Swanson, 2015)* |
| BMAa | *basomedial amygdalar nucleus anterior part (>1840)* |
| BMAp | *basomedial amygdalar nucleus posterior part (>1840)* |
| BST | *bed nuclei of terminal stria (Gurdjian, 1925)* |
| BSTal | *bed nuclei of terminal stria anterolateral area (Swanson, 2004)* |
| BSTam | *bed nuclei of terminal stria anteromedial area (Dong & Swanson, 2006)* |
| BSTfu | *bed nuclei of terminal stria fusiform nucleus (Ju & Swanson, 1989)* |
| BSTov | *bed nuclei of terminal stria anteromedial area (Ju & Swanson, 1989)* |
| CA1 | *Ammon's horn field CA1 (Lorente de Nó, 1934)* |
| CA1d | *Ammon's horn field CA1 dorsal part (Risold & Swanson, 1996)* |
| CA1v | *Ammon's horn field CA1 ventral part (Risold & Swanson, 1996)* |
| CA2 | *Ammon's horn field CA2 (Lorente de Nó, 1934)* |
| CA3so | *Ammon's horn field CA3 stratum oriens (>1840)* |
| CEA | *central amygdalar nucleus (Johnston, 1923)* |
| CEAc | *central amygdalar nucleus capsular part (McDonald, 1982)* |
| CEAl | *central amygdalar nucleus lateral part (Swanson, 1992)* |
| CEAm | *central amygdalar nucleus medial part (McDonald, 1982)* |
| CLA | *claustrum (Burdach, 1822)* |
| CLl | *central linear raphe nucleus (>1840)* |
| CM | *central medial thalamic nucleus (Rioch, 1929)* |
| COA | *cortical amygdalar area (>1840)* |
| COAa | *cortical amygdalar area anterior part (>1840)* |
| COApl | *cortical amygdalar area posterior part lateral zone (>1840)* |
| COApm | *cortical amygdalar area posterior part medial zone (>1840)* |
| CP | *caudoputamen (Heimer, 1975)* |
| cpd | *cerebral peduncle (Tarin, 1753)* |
| cVP | central aspect of ventral pallidum |
| DG | *dentate gyrus (>1840)* |
| DGmo | *dentate gyrus molecular layer (>1840)* |
| DGsg | *dentate gyrus granule cell layer (>1840)* |
| DL | dorsal striatum dorsolateral part |
| DM | dorsal striatum dorsomedial part |
| dmACB | dorsomedial column of accumbens nucleus |
| DP | dorsal peduncular cortex |
| DS | dorsal striatum |
| ec | *external capsule (Burdach, 1822)* |
| ECT | *ectorhinal area (Brodmann, 1909)* |
| ENTl | *entorhinal area lateral part (>1840)* |
| ENTm | *entorhinal area medial part (>1840)* |
| EPd | *endopiriform nucleus dorsal part (Krettek & Price, 1977)* |
| EPv | *endopiriform nucleus ventral part (Krettek & Price, 1977)* |
| frf | *radiation of corpus callosum frontal forceps (>1840)* |
| GPl | *lateral globus pallidus (>1840)* |
| GPm | *medial globus pallidus (>1840)* |
| HF | *hippocampal formation (Swanson & Cowan, 1977)* |
| I | *internuclear hypothalamic area (Swanson, 2004)* |
| IA | *intercalated amygdalar nuclei (>1840)* |
| IAD | *interanterodorsal thalamic nucleus (>1840)* |
| IF | *interfascicular raphe nucleus (>1840)* |
| ILA | *infralimbic area (Rose & Woolsey, 1948)* |
| int | *internal capsule (Burdach, 1822)* |
| IPN | *interpeduncular nucleus (>1840)* |
| isl | *islands of Calleja (>1840)* |
| LA | *lateral amygdalar nucleus (>1840)* |
| LHA | *lateral hypothalamic area (Nissl, 1913)* |
| LHAa | *lateral hypothalamic area anterior group anterior region (Swanson, 2004)* |
| LHAai | *lateral hypothalamic area anterior group anterior region intermediate zone (Swanson, 2004)* |
| LHAav | *lateral hypothalamic area anterior group anterior region ventral zone (Swanson, 2004)* |
| LHAjp | *lateral hypothalamic area middle group medial tier juxtaparaventricular region (Swanson, 2004)* |
| LP | *lateral posterior thalamic nucleus (>1840)* |
| LPO | *lateral preoptic nucleus (>1840)* |
| LS | *lateral septal nucleus (Ramón y Cajal, 1909–1911)* |
| LS.vl | *lateral septal nucleus, ventrolateral zone (Ramón y Cajal, 1909–1911)* |
| MD | *mediodorsal thalamic nucleus (>1840)* |
| MDm | *mediodorsal thalamic nucleus medial part (>1840)* |
| MEA | *medial amygdalar nucleus (Johnston, 1923)* |
| MEAad | *medial amygdalar nucleus anterodorsal part (>1840)* |
| MEApd | *medial amygdalar nucleus posterodorsal part (>1840)* |
| MEApd.a | *medial amygdalar nucleus posterodorsal part sublayer a (>1840)* |
| MEApd.b | *medial amygdalar nucleus posterodorsal part sublayer b (>1840)* |
| MEApd.c | *medial amygdalar nucleus posterodorsal part sublayer c (>1840)* |
| MEApv | *medial amygdalar nucleus posteroventral part (>1840)* |
| MGd | *medial geniculate complex dorsal part (>1840)* |
| MGv | *medial geniculate complex ventral part (>1840)* |
| MOs | *secondary somatomotor areas (>1840)* |
| mPFC | medial prefrontal cortex |
| MS | *medial septal nucleus (>1840)* |
| mVP | medial aspect of ventral pallidum |
| MRNm | *midbrain reticular nucleus (>1840)* |
| NB | *nucleus of brachium of inferior colliculus (>1840)* |
| NC | *nucleus circularis (>1840)* |
| NDB | *diagonal band nucleus (>1840)* |
| opth | *hypothalamic optic tract (Swanson, 2015)* |
| optt | *thalamic optic tract (Swanson, 2015)* |
| ORBl | *lateral orbital area (Krettek & Price, 1977)* |
| ORBm | *medial orbital area (Krettek & Price, 1977)* |
| ORBv | *ventral orbital area (Krettek & Price, 1977)* |
| ORBvl | *ventrolateral orbital area (Krettek & Price, 1977)* |
| OT | *olfactory tubercle (Calleja, 1893)* |
| PA | *posterior amygdalar nucleus (Canteras et al., 1992)* |
| PAA | *piriform-amygdalar area (Canteras et al., 1992)* |
| PAR | *parasubiculum (>1840)* |
| p-dmACB | dorsomedial column of accumbens nucleus, posterior part |
| PERI | *perirhinal area (Brodmann, 1909)* |
| PIR | *piriform area (Smith, 1919)* |
| PL | *prelimbic area (Brodmann, 1909)* |
| PMv | *ventral premammillary nucleus (>1840)* |
| pofh | *hypothalamic postcommissural fornix (Swanson, 2015)* |
| PP | *peripeduncular nucleus (>1840)* |
| PRE | *presubiculum (Swanson & Cowan, 1977)* |
| PS | *parastrial nucleus (Simerly et al., 1984)* |
| PT | *paratenial nucleus (>1840)* |
| PVa | *periventricular hypothalamic nucleus anterior part (Swanson, 2018)* |
| PVH | *paraventricular hypothalamic nucleus (>1840)* |
| PVHdp | *paraventricular hypothalamic nucleus descending division dorsal parvicellular part (>1840)* |
| PVHmpd | *paraventricular hypothalamic nucleus parvicellular division medial parvicellular part dorsal zone (Simmons & Swanson, 2008)* |
| PVHpmm | *paraventricular hypothalamic nucleus magnocellular division posterior magnocellular part medial zone (>1840)* |
| PVHpv | *paraventricular hypothalamic nucleus parvicellular division periventricular part (>1840)* |
| PVT | *paraventricular thalamic nucleus (>1840)* |
| RCH | *lateral hypothalamic area anterior group retrochiasmatic area (>1840)* |
| rhs | *rhinal sulcus (>1840)* |
| RT | *reticular thalamic nucleus (>1840)* |
| SBPV | *subparaventricular zone (Watts et al., 1987)* |
| SF | *septofimbrial nucleus (>1840)* |
| SGN | *suprageniculate nucleus (>1840)* |
| SI | *substantia innominata* or *innominate substance (Schwalbe, 1881)* |
| smh | *hypothalamic medullary stria (Swanson, 2018)* |
| SNc | *substantia nigra compact part (Sano, 1910)* |
| SNr | *substantia nigra reticular part (Sano, 1910)* |
| SOp | *supraoptic nucleus principal part (Swanson, 2018)* |
| SPFpl | *subparafascicular nucleus parvicellular part lateral division (>1840)* |
| ste | *endbrain terminal stria (Swanson, 2015)* |
| STN | *subthalamic nucleus (>1840)* |
| SUB | *subiculum (>1840)* |
| SUBd | *subiculum dorsal part (Swanson & Cowan, 1975)* |
| SUBv | *subiculum ventral part (Swanson & Cowan, 1975)* |
| suph | *hypothalamic supraoptic decussations (Swanson, 2018)* |
| TEa | *temporal association areas (Swanson, 1992)* |
| TEP | *temporal pole (Broca, 1878)* |
| TR | *postpiriform transition area (Canteras et al., 1992)* |
| TTd | *tenia tecta dorsal part (Swanson, 1992)* |
| TUte | *terete part of the tuberal nucleus (Petrovich et al., 2001)* |
| VL | dorsal striatum ventrolateral part |
| vlACB | ventrolateral aspect of accumbens nucleus |
| VM | dorsal striatum ventromedial part |
| vmACB | ventromedial aspect of accumbens nucleus |
| VLi | *inferior horn of lateral ventricle (Bell, 1802)* |
| vlt | *ventrolateral hypothalamic tract (Swanson, 2004)* |
| VMH | *ventromedial hypothalamic nucleus (>1840)* |
| VP | ventral pallidum |
| VTA | *ventral tegmental area (Tsai, 1925)* |
| ZI | *zona incerta (>1840)* |

We derived standard terms for gray matter regions from “Table C: Rat CNS Gray Matter Regions 4.0 (Topographical Histological Groupings)” in the Supporting Information for Swanson (Swanson, 2018), which is based on the literature and cytoarchitectonic features of Nissl-stained sections. We include all citations embedded within standard terms in the reference list. We also provide a complete list of standard terms and abbreviations used in our study **(Table 1)**.

In addition, we use the abbreviations “dmACB” and “aPVT,” which are not formally included in Swanson (2018) but follow its abbreviation system and its extension in Thompson and Swanson (2010). We also use terms for regions not yet formalized in *Brain Maps* at the time of writing, such as VP (ventral pallidum) and DP (dorsal peduncular cortex). Future atlas versions will need to formally incorporate or refine these designations.

We place non-standard lowercase modifiers (not part of the *Brain Maps* system) before the main abbreviation (e.g., aPVT, mVP), whereas standard lowercase modifiers follow the main capitalized abbreviation, consistent with *Brain Maps* conventions (e.g., LHAa).

Finally, the ACB is commonly recognized by its core-shell distinction and alternative nomenclature (i.e. “NAc shell” and “NAc core”). We retained the Swanson nomenclature system, but we included calbindin staining as a classical marker for delineating core-shell boundaries (Voorn et al., 1989) within the *Brain Maps 4.0* spatial framework. The more familiar ACB core and shell divisions outlined in Paxinos and Watson (2014) were based mainly on cholecystokinin and substance P immunoreactivity, but several reports recognized calbindin as a robust alternative (Jongen-Rêlo et al., 1994; Zahm & Brog, 1992). Using this marker, we can approximate our dmACB as ‘dorsomedial ACB shell.’

### Experiment 2: dmACB neuronal activation during incubation of oxycodone seeking

#### Subjects

In Experiment 2, we used male (n=14) and female (n=14) rats from the same source and weighing 190–275 g prior to surgery. We maintained rats under a reverse 12:12 h light/dark cycle (lights off at 8:00 A.M.) with free access to food and water. We housed two rats per cage prior to surgery and individually after surgery. We performed Experiment 2 in accordance with the NIH *Guide for the Care and Use of Laboratory Animals (8th edition)*, under protocols approved by the Animal Care and Use Committee of NIDA Intramural Research Program.

#### Intravenous surgery

We anesthetized the rats with isoflurane (5% induction, 2–3% maintenance, Covetrus). We attached silastic catheters to a modified 22-gauge cannula cemented to polypropylene mesh (Amazon or Industrial Netting), inserted the catheter into the jugular vein and fixed the mesh to the mid-scapular region of the rat (Caprioli et al., 2018; Caprioli et al., 2017; Pitts et al., 2026; Venniro et al., 2018). We injected the rats with ketoprofen (2.5 mg/kg, s.c., Covetrus) after surgery and the following day to relieve pain and decrease inflammation. We allowed the rats to recover for 6–8 days before oxycodone self-administration training. During recovery and all experimental phases, we flushed the catheters every 24–48 h with gentamicin (4.25–5.0 mg/ml, Fresenius Kabi or Covetrus) dissolved in sterile saline. If we suspected catheter failure during training, we tested patency with Diprivan (propofol, NIDA pharmacy, 10 mg/mL, 0.1–0.2 mL injection volume, i.v.), and if not patent, we re-catheterized the other jugular vein and continued training the next day or eliminated the rat from the study.

#### Drugs

We received oxycodone hydrochloride (HCl) from NIDA pharmacy and dissolved it in sterile saline. We chose a unit dose of 0.1 mg/kg for self-administration training based on our previous studies (Fredriksson et al., 2020; Fredriksson et al., 2023; Negishi et al., 2025).

#### Oxycodone self-administration training

We trained the rats to self-administer oxycodone-HCl for 6-h/day for 14 days under the FR1-20-s timeout reinforcement schedule. Oxycodone was infused at a volume of 100 µl over 3.5 s at a unit dose of 0.1 mg/kg/infusion. Each session began with illumination of a red house light that remained on for the entire session, followed 10-s later by the insertion of the active lever. Active lever presses led to oxycodone infusions that were paired with a 3.5-s tone-light cue. We limited oxycodone intake to 100 infusions per session.

#### Electric barrier phase

During this phase, oxycodone was available for 2-h per day for 10–11 days. We used the same parameters (oxycodone dose, reinforcement schedule, tone-light cues, etc.) that we used during the training phase. To achieve abstinence, we introduced an electric barrier near the active lever (“shock zone”) (Cooper et al., 2007; Fredriksson et al., 2020). We separated the “shock zone” (2/3 of the chamber) from a “safe zone” (remaining 1/3 of the chamber) with a plastic demarcation (McMaster, cat # 9852K61). If the rats approached the active lever, they received a continuous mild footshock (0.1–0.4 mA). On the first day, we set the current at 0.0 mA and then gradually increased the intensity to 0.3 mA (0.1 mA increments per day). If a rat did not suppress oxycodone self-administration (< 3 infusions per day), we increased the intensity to 0.4 mA the next day. Prior to the electric barrier phase, we tested the rats’ sensitivity to footshock (operationally defined as the minimal shock level that causes the withdrawal of the front paw). There were no group or sex differences in shock sensitivity, assessed with 0.05 mA increments, starting at 0.05 mA, and the values for individual rats ranged from 0.1 mA to 0.2 mA.

#### Relapse (incubation) tests

We tested the rats for oxycodone seeking under extinction conditions during early abstinence (day 1, before electric barrier exposure) or late abstinence (day 15, after electric barrier exposure). Before testing, we placed all rats in the self-administration chamber for a 30-min habituation period to ensure that the day 15 rats recognized that the barrier was no longer electrified. We conducted a 120-min relapse test. During the test, active lever presses delivered the oxycodone-paired compound tone-light cue and activated the infusion pump but did not deliver drug. We turned off the electric barrier during testing for the day 15 rats. For Fos analysis, we perfused some rats immediately after the relapse test on day 1 (n = 9) or day 15 (n = 9). We perfused additional rats immediately after removing them from their home cage on day 2 (n = 5) or day 16 (n = 5); these rats served as the no-test controls.

#### Fos detection and quantification

We followed the IHC and microscopy procedures described above with some differences for Fos detection. For every step of tissue processing, we included rats from all four groups (day 1 no-test, day 1 test, day 15 no-test, day 15 test) to mitigate systematic differences among the groups. We used an anti-phospho-c-Fos (Ser^32^) monoclonal antibody (host: Rabbit; dilution: 1:4,000; Cell Signaling Technology; Cat# 5348; RRID: AB_10557109) and anti-calbindin (host: Mouse; dilution: 1:4,000; Millipore-Sigma; Cat# C9848; RRID: AB_476894). The anti-phospho-c-Fos antibody we used recognizes Fos only when phosphorylated at serine 32 and does not cross-react with FosB, FRA1, or FRA2. Accordingly, throughout this study “Fos” refers to pSer^32^-Fos immunoreactivity (pSer^32^-Fos-ir) in our material, which we used as a proxy for Fos induction, as in prior studies of incubation of drug craving (Caprioli et al. 2017); Ser^32^ phosphorylation has been reported to stabilize Fos protein (Sasaki et al. 2006). To our knowledge, the correspondence between pSer^32^-Fos-ir and total Fos immunoreactivity has not been directly established in rat brain, and we therefore interpret Fos-labeled cell counts as an index of neuronal activation with this caveat. The secondary antibody cocktail contained AlexaFluor 555-conjugated antibody raised against rabbit (host: Donkey; dilution: 1:500; Jackson ImmunoResearch Labs; Cat# 711-565-152; RRID: AB_3095471), AlexaFluor 488-conjugated antibody raised against mouse (host: Donkey; dilution: 1:500; Jackson ImmunoResearch Labs; Cat# 715-545-151; RRID: AB_2341099) and the counterstain we used was NeuroTrace 660/640 (Thermo Fisher Scientific).

We obtained all images in a single slide scanning session using the same imaging parameters and exported channels separately as 16-bit grayscale images. After consulting histograms from several sections, we applied the same image rescaling parameters to each image. We used a Cellpose*-*SAM model (Stringer & Pachitariu, 2025) trained to identify Fos-positive nuclei to generate segmentations for each image. To assess the robustness of image scaling and Fos detection across experimental groups, we performed histogram-based analyses of signal intensity distributions.

For segmented objects, we computed the mean pixel intensity within each Fos-positive nucleus, yielding a distribution of per-object Fos intensities. To estimate background signal, we sampled all non-zero image pixels excluding those within Fos segmentations, generating a complementary distribution of background intensities. Histograms for both Fos and background signals were computed using 4-unit bins across the 0–255 intensity range. These distributions were used to quantify signal-to-noise separation using a signal-to-noise ratio (SNR) metric defined as:

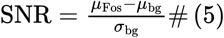

Here, *μ_Fos_* and *μ_bg_* represent the mean Fos and background intensities, respectively, and σ_bg_ is the standard deviation of the background distribution. This metric quantifies the degree to which segmented Fos signals are elevated above background noise, allowing comparison of segmentation performance and signal separation across experimental conditions.

After aligning Fos segmentations to their original images using Adobe Illustrator, we included boundary ROIs for anterior and posterior ACB subregions. We drew ROIs based on classical core and shell divisions revealed using calbindin staining (Jongen-Rêlo et al., 1994), and we approximated dorsomedial and ventrolateral shell boundaries using the dmACB locations identified in Exp. 1. We exported Fos segmentations and ROIs as SVG files and used a custom Python script to measure ROI areas and respective Fos counts. Fos-positive cells were counted blindly and independently by two people (KN and MSP; inter-rater reliability, Pearson *r* = 0.997).

#### Statistical analysis

We analyzed behavioral and Fos immunohistochemistry data using repeated-measures ANOVAs (RM-ANOVAs) or ANOVAs in SPSS (Version 31, GLM procedure). When main effects or interactions were significant (p < 0.05), we conducted followed-up analyses using one-way ANOVAs and/or Tukey’s tests. When sphericity assumptions were violated, we applied the Greenhouse-Geisser correction. We describe the specific between- and within-subject factors for each analysis in the Results section. Because the multifactorial ANCOVAs yielded multiple main and interaction effects, we report only those that are critical for data interpretation. For clarity, we indicate post-hoc results with asterisks in the Figures but do not describe them in the Results section.

We included sex as a factor in the behavioral analyses (n = 14 per sex during self-administration training and electric barrier exposure; n = 7 per sex for relapse tests on abstinence days 1 and 15). We did not include Sex in the Fos analyses because the number of males and females per condition (n = 4–5) was too small to support reliable statistical comparisons. For Fos, we analyzed three groups: Day 1 test (n = 9, 5 females), Day 15 test (n = 9, 4 females), and No-Test (n = 10, 5 females; see Results).

Inactive lever presses, a measure of non-specific activity and/or response generalization (Shalev et al., 2002) were very low across all phases of the experiment (see **Fig. 13**). Therefore, we included this measure as a covariate in the statistical analyses. We provide complete statistical results for **Figures 13 and 14** in **Table 2**.

**Table 2.** Summary of statistical results described in. **Figures 13 and 14**.

| Figure number | Factor name | F-value | *p*-value | Partial Eta<sup>2</sup> |
|---|---|---|---|---|
| **Figure 13B**. Self-administration: Infusions | Session within-subjects<br>Sex (male, female) between-subjects<br>Session × Sex | F<sub>(3.6, 94.1)</sub> = 26.4<br>F<sub>(1, 26)</sub> = 18.1<br>F<sub>(3.6, 94.1)</sub> = 5.9 | <0.001<br><0.001<br><0.001 | 0.504<br>0.410<br>0.185 |
| **Figure 13B**. Self-administration:<br>Active lever presses | Session within-subjects<br>Sex (male, female) between-subjects<br>Session × Sex<br>Inactive lever: covariate | F<sub>(1.7,42.0)</sub> = 4.9<br>F<sub>(1,25)</sub> = 8.1<br>F<sub>(1.7, 42.0)</sub> = 4.0 | 0.017<br>0.009<br>0.031 | 0.163<br>0.245<br>0.139 |
| **Figure 13C**. Electric barrier: Infusions | Session within-subjects<br>Sex between-subjects<br>Session × Sex | F<sub>(10,120)</sub> = 125.3<br>F<sub>(1,12)</sub> = 19.5<br>F<sub>(10,120)</sub> = 15.0 | <0.001<br><0.001<br><0.001 | 0.904<br>0.619<br>0.554 |
| **Figure 13C**. Electric barrier:<br>Active lever presses | Session within-subjects<br>Sex between-subjects<br>Session × Sex<br>Inactive lever: covariate | F<sub>(10,110)</sub> = 14.3<br>F<sub>(1,11)</sub> = 5.9<br>F<sub>(10,110)</sub> = 4.9 | <0.001<br>0.034<br><0.001 | 0.566<br>0.348<br>0.307 |
| **Figure 13D**. Relapse tests total<br>Active Lever presses | Abstinence day between-subjects<br>Sex between subjects<br>Abstinence day × Sex<br>Inactive lever: covariate | F<sub>(1,23)</sub> = 32.0<br>F<sub>(1,23)</sub> = 9.9<br>F<sub>(1,23)</sub> = 3.4 | <0.001<br>0.005<br>0.079 | 0.582<br>0.300<br>0.128 |
| **Figure 13D**. Relapse tests time course<br>Active lever presses | Abstinence day between-subjects<br>Session time within-subjects<br>Sex between-subjects<br>Abstinence day × Session time<br>Abstinence day × Sex<br>Session time × Sex<br>Abstinence day × Session time × Sex<br>Inactive lever: covariate | F<sub>(1,23)</sub> = 32.0<br>F<sub>(1.3,30.6)</sub> = 34.2<br>F<sub>(1,23)</sub> = 9.9<br>F<sub>(1.3, 30.6)</sub> = 11.7<br>F<sub>(1,23)</sub> = 3.4<br>F<sub>(1.3, 30.6)</sub> = 6.2<br>F<sub>(1.3, 30.6)</sub> = 1.3 | <0.001<br><0.001<br>0.005<br><0.001<br>0.079<br>0.012<br>0.272 | 0.582<br>0.598<br>0.300<br>0.338<br>0.128<br>0.211<br>0.054 |
| **Figure 12F**. Segmentation Evaluation of SNR | Test condition between-subjects | F<sub>(2,25)</sub> = 1.1 | 0.352 | 0.080 |
| **Figures 14B & 14C**. Overall Analysis Fos/mm<sup>2</sup> | Striatal region within-subjects<br>Bregma level within-subjects<br>Test condition between-subjects<br>Region × Test condition<br>Bregma × Test condition<br>Region × Bregma<br>Region × Bregma × Test condition | F<sub>(2.2,54.1)</sub> = 117.0<br>F<sub>(1, 25)</sub> = 23.6<br>F<sub>(2,25)</sub> = 44.2<br>F<sub>(4.3,54.1)</sub> = 15.7<br>F<sub>(2,25)</sub> = 9.1<br>F<sub>(3.2, 80.2)</sub> = 19.6<br>F<sub>(6.4, 80.2)</sub> = 6.4 | <0.001<br><0.001<br><0.001<br><0.001<br>0.001<br><0.001<br><0.001 | 0.824<br>0.485<br>0.779<br>0.556<br>0.422<br>0.440<br>0.338 |
| **Figure 14B.**<br>Fos/mm<sup>2</sup> in anterior ACBcore | Test condition between-subjects<br>Tukey post-hoc test<br>No test vs Day 1<br>No test vs Day 15<br>Day 1 vs Day 15 | F<sub>(2,25)</sub> = 27.9 | <0.001<br>--------<br><0.001<br><0.001<br>0.078 | 0.691 |
| **Figure 14B.**<br>Fos/mm<sup>2</sup> in posterior ACBcore | Test condition) between-subjects<br>Tukey post-hoc test<br>No test vs Day 1<br>No test vs Day 15<br>Day 1 vs Day 15 | F<sub>(2,25)</sub> = 23.5 | <0.001<br>--------<br><0.001<br><0.001<br>0.339 | 0.653 |
| **Figure 14B.**<br>Fos/mm<sup>2</sup> in anterior dmACB | Test condition between-subjects<br>Tukey post-hoc test<br>No test vs Day 1<br>No test vs Day 15<br>Day 1 vs Day 15 | F<sub>(2,25)</sub> = 29.5 | <0.001<br>--------<br><0.001<br>0.002<br>**0.003** | 0.702 |
| **Figure 14B.**<br>Fos/mm<sup>2</sup> in posterior dmACB | Test condition) between-subjects<br>Tukey post-hoc test<br>No test vs Day 1<br>No test vs Day 15<br>Day 1 vs Day 15 | F<sub>(2,25)</sub> = 29.8 | <0.001<br>--------<br><0.001<br><0.001<br>**0.050** | 0.705 |
| **Figure 14B.**<br>Fos/mm<sup>2</sup> in anterior vmACB | Test condition between-subjects<br>Tukey post-hoc test<br>No test vs Day 1<br>No test vs Day 15<br>Day 1 vs Day 15 | F<sub>(2,25)</sub> = 31.3 | <0.001<br>--------<br><0.001<br><0.001<br>0.123 | 0.714 |
| **Figure 14B.**<br>Fos/mm<sup>2</sup> in posterior vmACB | Test condition) between-subjects | F<sub>(2,25)</sub> = 2.4 | 0.107 | 0.163 |
| **Figure 14B.**<br>Fos/mm<sup>2</sup> in anterior vlACB | Test condition between-subjects<br>Tukey post-hoc test<br>No test vs Day 1<br>No test vs Day 15<br>Day 1 vs Day 15 | F<sub>(2,25)</sub> = 17.8 | <0.001<br>--------<br><0.001<br>0.003<br>0.112 | 0.588 |
| **Figure 14B.**<br>Fos/mm<sup>2</sup> in posterior vlACB | Test condition) between-subjects<br>Tukey post-hoc test<br>No test vs Day 1<br>No test vs Day 15<br>Day 1 vs Day 15 | F<sub>(2,25)</sub> = 26.4 | <0.001<br>--------<br><0.001<br><0.001<br>**0.019** | 0.678 |
| **Figure 14C.**<br>Fos/mm<sup>2</sup> in anterior Dorsomedial striatum | Test condition between-subjects<br>Tukey post-hoc test<br>No test vs Day 1<br>No test vs Day 15<br>Day 1 vs Day 15 | F<sub>(2,25)</sub> = 15.9 | <0.001<br>--------<br><0.001<br><0.001<br>0.499 | 0.560 |
| **Figure 14C.**<br>Fos/mm<sup>2</sup> in posterior Dorsomedial striatum | Test condition) between-subjects<br>Tukey post-hoc test<br>No test vs Day 1<br>No test vs Day 15<br>Day 1 vs Day 15 | F<sub>(2,25)</sub> = 19.0 | <0.001<br>--------<br><0.001<br><0.001<br>0.685 | 0.603 |
| **Figure 14C.**<br>Fos/mm<sup>2</sup> in anterior Dorsolateral striatum | Test condition between-subjects<br>Tukey post-hoc test<br>No test vs Day 1<br>No test vs Day 15<br>Day 1 vs Day 15 | F<sub>(2,25)</sub> = 17.1 | <0.001<br>--------<br><0.001<br>0.075<br>**0.006** | 0.578 |
| **Figure 14C.**<br>Fos/mm<sup>2</sup> in posterior Dorsolateral striatum | Test condition) between-subjects<br>Tukey post-hoc test<br>No test vs Day 1<br>No test vs Day 15<br>Day 1 vs Day 15 | F<sub>(2,25)</sub> = 10.3 | <0.001<br>--------<br><0.001<br>0.023<br>0.267 | 0.452 |
| **Figure 14C.**<br>Fos/mm<sup>2</sup> in anterior Ventromedial dorsal striatum | Test condition between-subjects<br>Tukey post-hoc test<br>No test vs Day 1<br>No test vs Day 15<br>Day 1 vs Day 15 | F<sub>(2,25)</sub> = 19.1 | <0.001<br>--------<br><0.001<br><0.001<br>0.911 | 0.605 |
| **Figure 14C.**<br>Fos/mm<sup>2</sup> in posterior Ventromedial dorsal striatum | Test condition) between-subjects<br>Tukey post-hoc test<br>No test vs Day 1<br>No test vs Day 15<br>Day 1 vs Day 15 | F<sub>(2,25)</sub> = 16.5 | <0.001<br>--------<br><0.001<br>0.002<br>0.269 | 0.569 |
| **Figure 14C.**<br>Fos/mm<sup>2</sup> in anterior Ventrolateral dorsal striatum | Test condition between-subjects<br>Tukey post-hoc test<br>No test vs Day 1<br>No test vs Day 15<br>Day 1 vs Day 15 | F<sub>(2,25)</sub> = 18.0 | <0.001<br>--------<br><0.001<br>0.014<br>**0.022** | 0.591 |
| **Figure 14C.**<br>Fos/mm<sup>2</sup> in posterior Ventrolateral dorsal striatum | Test condition) between-subjects<br>Tukey post-hoc test<br>No test vs Day 1<br>No test vs Day 15<br>Day 1 vs Day 15 | F<sub>(2,25)</sub> = 14.2 | <0.001<br>--------<br><0.001<br>0.028<br>**0.048** | 0.532 |

## Results

### Experiment 1: Anatomical characterization of the dmACB column and circuit organization

#### Calbindin-based delineation of ACB

We used calbindin labeling to delineate ACB core and shell subdivisions across the anterior-posterior extent of ACB **(Fig. 1A)**. Our samples showed clear calbindin-poor zones in the ventral ACB that mark the shell subdivision. We then used a separate Neurotrace channel to register a representative ACB core-shell boundary to the *Brain Maps 4.0* atlas framework **(Fig. 1B)**. In the following experiments, we used either inferred atlas representations (Experiment 1) or empirically detected (Experiment 2) core-shell boundaries.is only found in the ventral zone of LHAa. The cytoarchitecture of paraventricular (PVH) and anterior (AHN) hypothalamic nuclei suggest this injection is at ^L25^AP ^β–1.5^. Based on its location in the LHAa ventral zone, this injection may also overlap with the retinorecipient zone of LHAa (Canteras et al., 2011).

**Figure 1.**
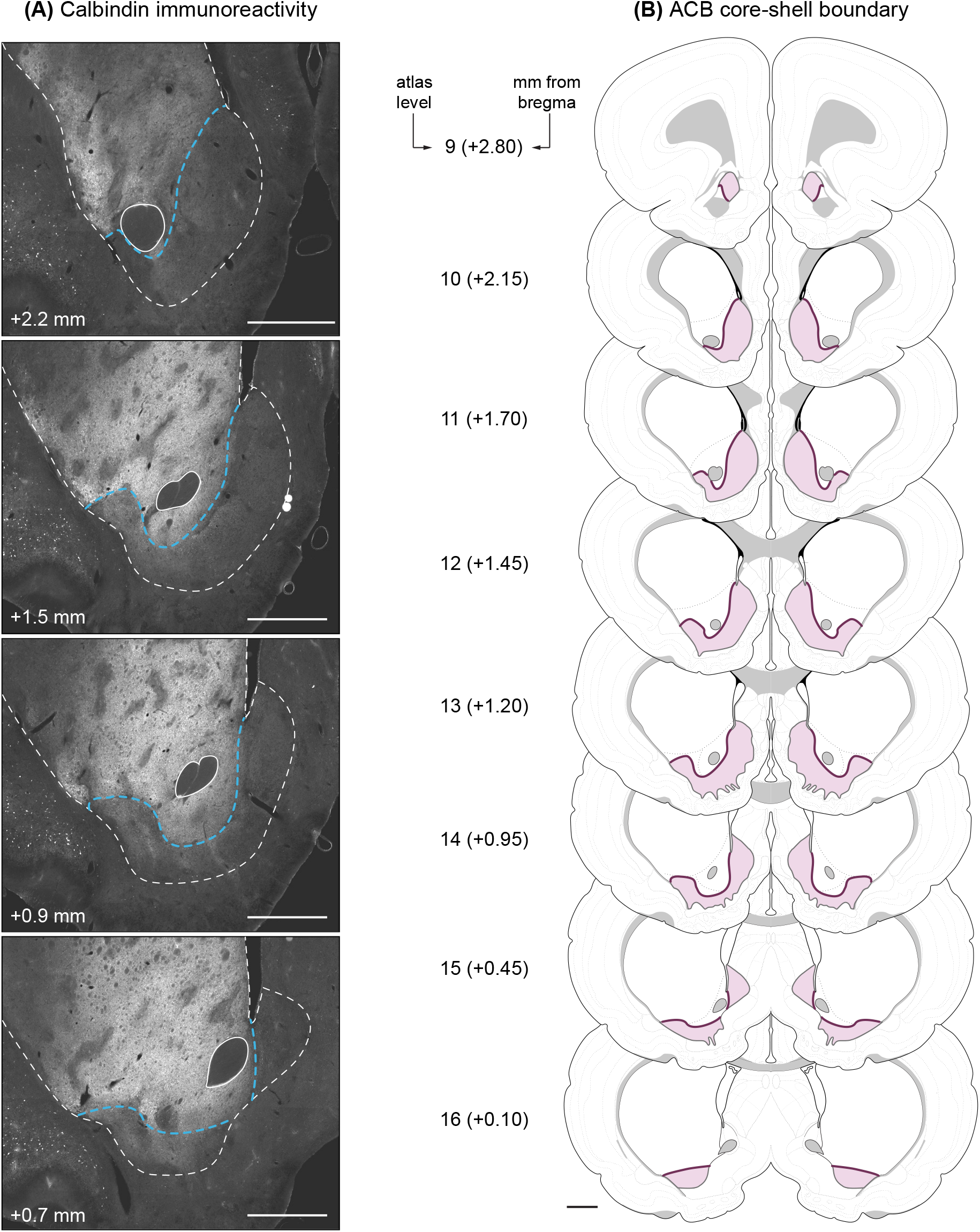
Atlas representation of the ACB core-shell boundary. **(A)** Photomicrographs showing calbindin immunoreactivity at different anterior-posterior positions of ACB. Dashed blue lines indicate the boundary interpretation. Dashed white lines indicate the striatal boundary determined using the NeuroTrace channel. **(B)** Atlas representations of core-shell boundary in purple. Shaded areas indicate ACB shell areas. mm values indicate approximate AP distance from Bregma. Scale bars: 1 mm **(A)**, 1 mm in atlas scale **(B)**.

#### Convergent aPVT, LHAa, and mVP connections in dmACB

Previous work has shown that PVT projections and retrograde labeling from LHAa overlap in ACB (Thompson & Swanson, 2010). In our initial experiments, we quantified the extent of overlap between PVT inputs and LHAa-projecting neurons throughout the ACB. To do this, we analyzed PHAL+CTB co-injection cases in which PHAL and CTB injection sites were restricted to aPVT **(Fig. 2A)** and LHAa **(Fig. 2B)**, respectively. We limited our analysis to brain sections that contained the ACB **(Fig. 2C)**.

**Figure 2.**
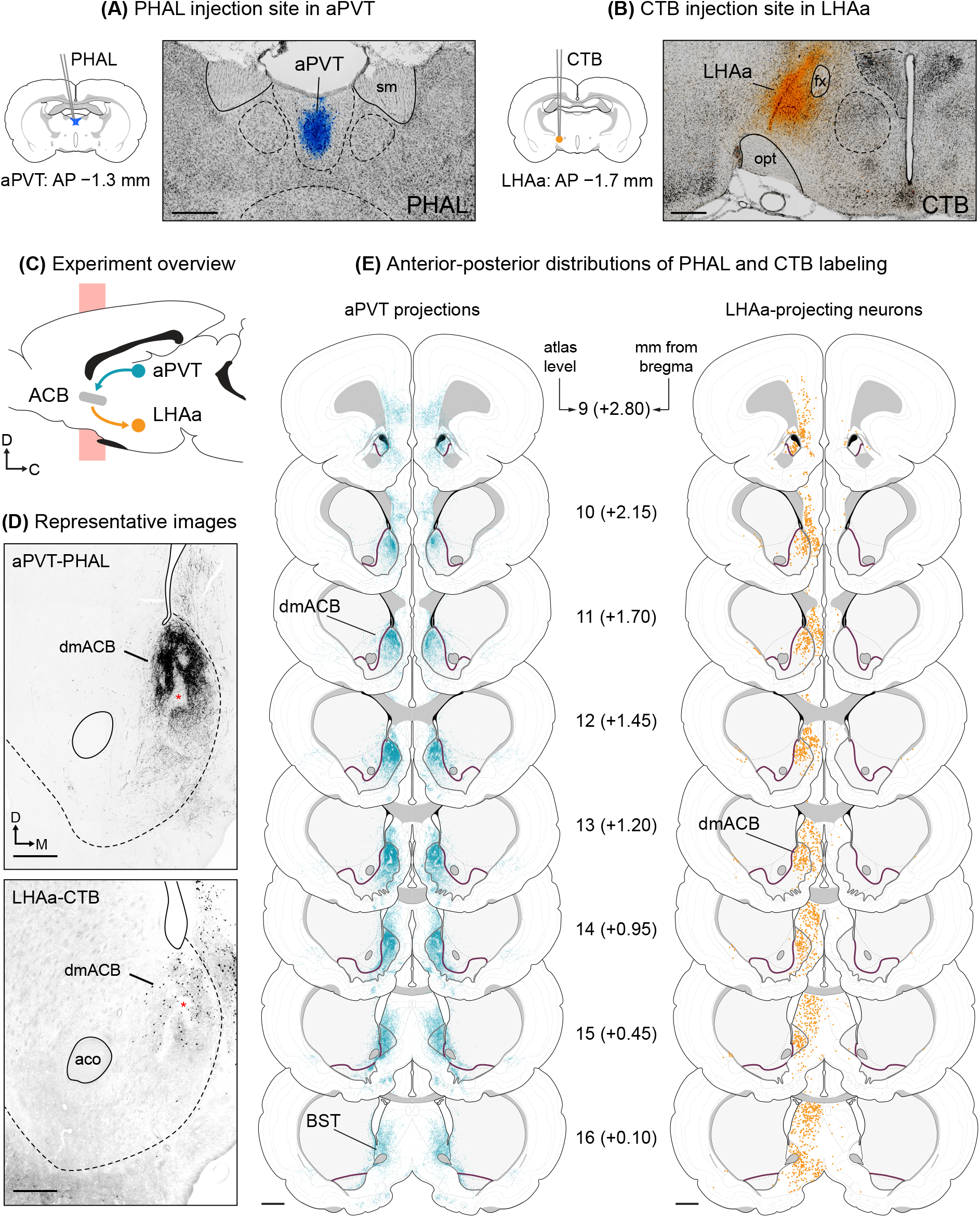
Mapping of the aPVT–dmACB–LHAa circuit motif. Photomicrographs showing NeuroTrace staining with pseudo-colored injection sites for **(A)** PHAL in aPVT and **(B)** CTB in LHAa. **(C)** Diagram of the pathway tracing experiment. **(D)** Epifluorescence photomicrographs showing (above) PHAL-labeled axons in dmACB following an injection into aPVT and (below) CTB-labeled neurons in dmACB following an injection into LHAa. Boundaries shown in **(A, B, and D)** were determined using the NeuroTrace channel. Asterisks in **(D)** indicate putative striosomal compartments. **(E)** Mapped distributions of aPVT axons (teal) and LHAa-projecting neurons (orange) across the anterior-posterior span of ACB. Dorsal striatum and ACB are indicated using solid boundaries and gray shading. Solid purple lines indicate the calbindin-based ACB core-shell boundaries (see Fig. 1). Values in **(E)** denote AP distance from bregma in mm. See **Table 1** for abbreviations. Scale bars: 500 µm **(A, B, D)**, 1 mm in atlas scale **(E)**.

We observed highly localized and dense aPVT axon terminals in a dorsomedial aspect of ACB **(Fig. 2D)**. The aPVT axons were almost absent within striosome-like dmACB spaces. Similarly, retrograde labeling from LHAa localized to the dmACB and was not observed in striosome-like spaces **(Fig. 2D)**. Full mapping of LHAa and aPVT connections in the ACB showed that both regions were specifically connected to the dmACB throughout its roughly 2-mm anterior-posterior span **(Fig. 2E)**. We observed aPVT axons and LHAa CTB labeling in the adjacent *bed nuclei of terminal stria (Gurdjian, 1925)* (BST), but labeling was lower compared to the dmACB.

The ACB forms a well-known projection to the VP that is topographically organized (Groenewegen et al., 1999). Therefore, we examined the distributions of CTB in the ACB after small injections into the mVP **(Fig. 3A)**. We selected two cases in which CTB deposit was restricted to a medial (mVP; alternatively, *innominate substance (Schwalbe, 1881)* (SI)) and a central aspect of VP (cVP) **(Fig. 3A)**. Our goal was to determine whether the VP medial-lateral axis is anatomically segregated into distinct striatopallidal projections.

**Figure 3.**
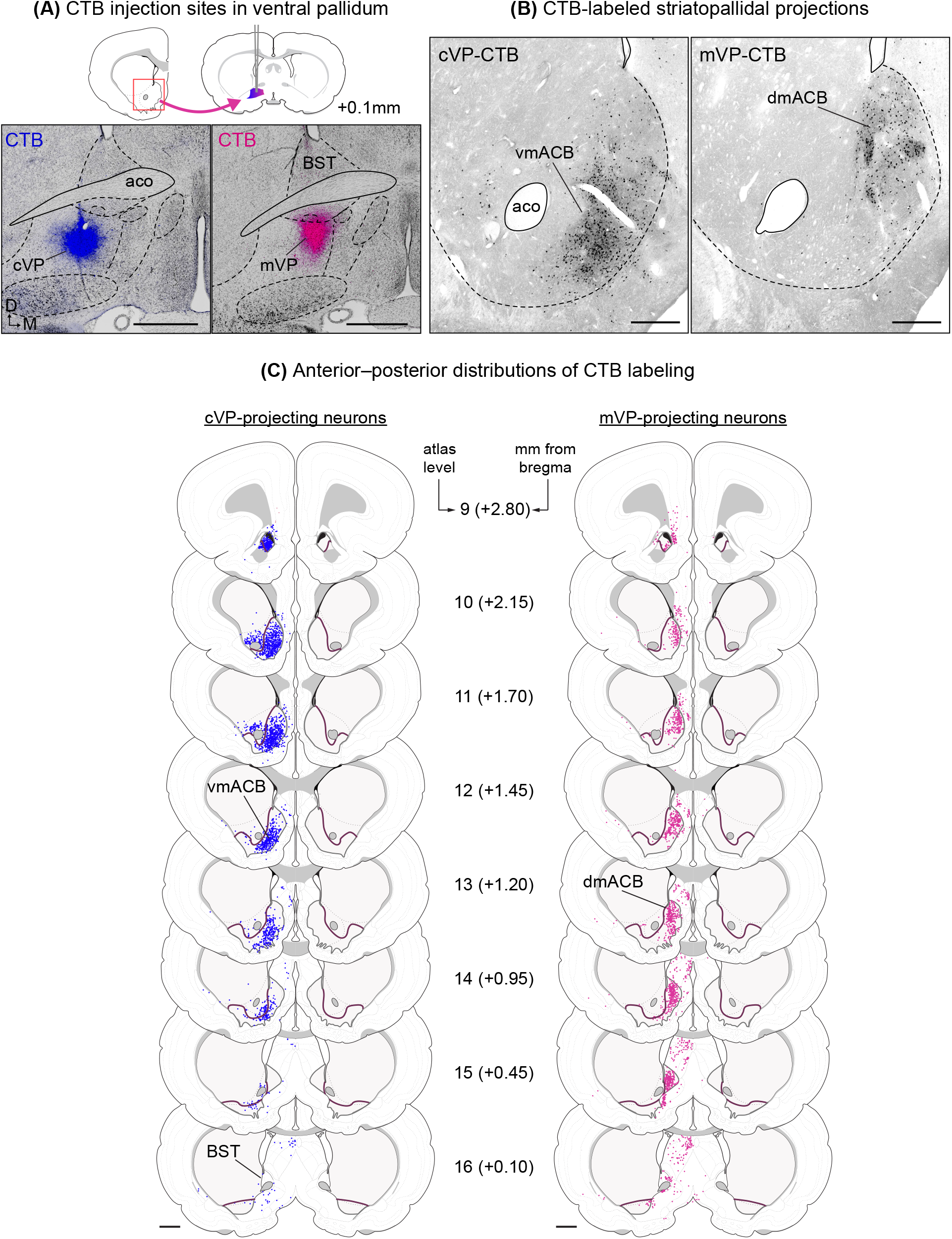
Retrograde tracing of striatopallidal projections. **(A)** *Above*: Atlas representation of CTB injection sites in VP. *Left*: Photomicrographs showing Neurotrace staining with pseudo-colored CTB injection sites in mVP in magenta. *Right*: Epifluorescence photomicrographs showing CTB-labeled neurons in dmACB following an injection into mVP. Boundaries were determined using the Neurotrace channel. **(B)** Mapped distributions of mVP-projecting neurons (magenta) across the anterior–posterior span of ACB. Dorsal striatum and ACB are indicated using solid boundaries and gray shading. Solid purple lines indicate the calbindin-defined ACB core-shell boundaries (see Fig. 1). Values to the left denote the atlas level in *Brain* Maps 4*.0* (Swanson, 2018), followed in parentheses by the approximate distance each level is from bregma in mm. See **Table 1** for abbreviations. Scale bars: 500 µm (**A**, right), 1 mm (**A**, left), 1 mm in atlas scale **(B)**.

We observed that tracer deposits restricted to the mVP and cVP labeled different parts of the ACB, respectively labeling dorsomedial and ventromedial subdomains **(Fig. 3B)**. This distribution pattern appears to be maintained across the anteroposterior axis of the ACB. There was no complete segregation between mVP and cVP distributions, but the injection sites we analyzed appear to occupy some common space in VP. Note that dmACB was consistently labeled with CTB from the mVP whereas CTB from the cVP sparsely distributes into the dmACB except in its most anterior part **(Fig. 3C)**. These results show that striatopallidal projections are spatially organized such that partly overlapping CTB injections in VP produce distinct distribution profiles in the ACB.

#### A connectional basis for defining the dmACB circuit

We showed that mVP, LHAa, and aPVT are preferentially connected with dmACB rather than adjacent striatal subdomains. To interpret these regions as components of a segregated basal ganglia loop, however, their connections must also be topographically co-spatial, occupying the same mesoscale anatomical domain. We devised intersection maps to objectively quantify and visualize the spatial convergence of connection densities. We first applied this approach to identify the mVP component of the circuit.

Previous work from Thompson and Swanson (2010) showed that dmACB-recipient mVP and LHAa subdomains project to aPVT. However, the corresponding indirect pathway linking dmACB to LHAa through mVP had not been anatomically localized. We therefore examined the spatial relationship between dmACB projections and LHAa-projecting neurons within VP using PHAL injections in dmACB and CTB injections in LHAa (**Fig. 4A**; see **Fig. 8** for injection sites). Labeling from these experiments showed prominent overlap within mVP, and VP labeling was sparse outside of this zone **(Fig. 4B)**.

**Figure 4.**
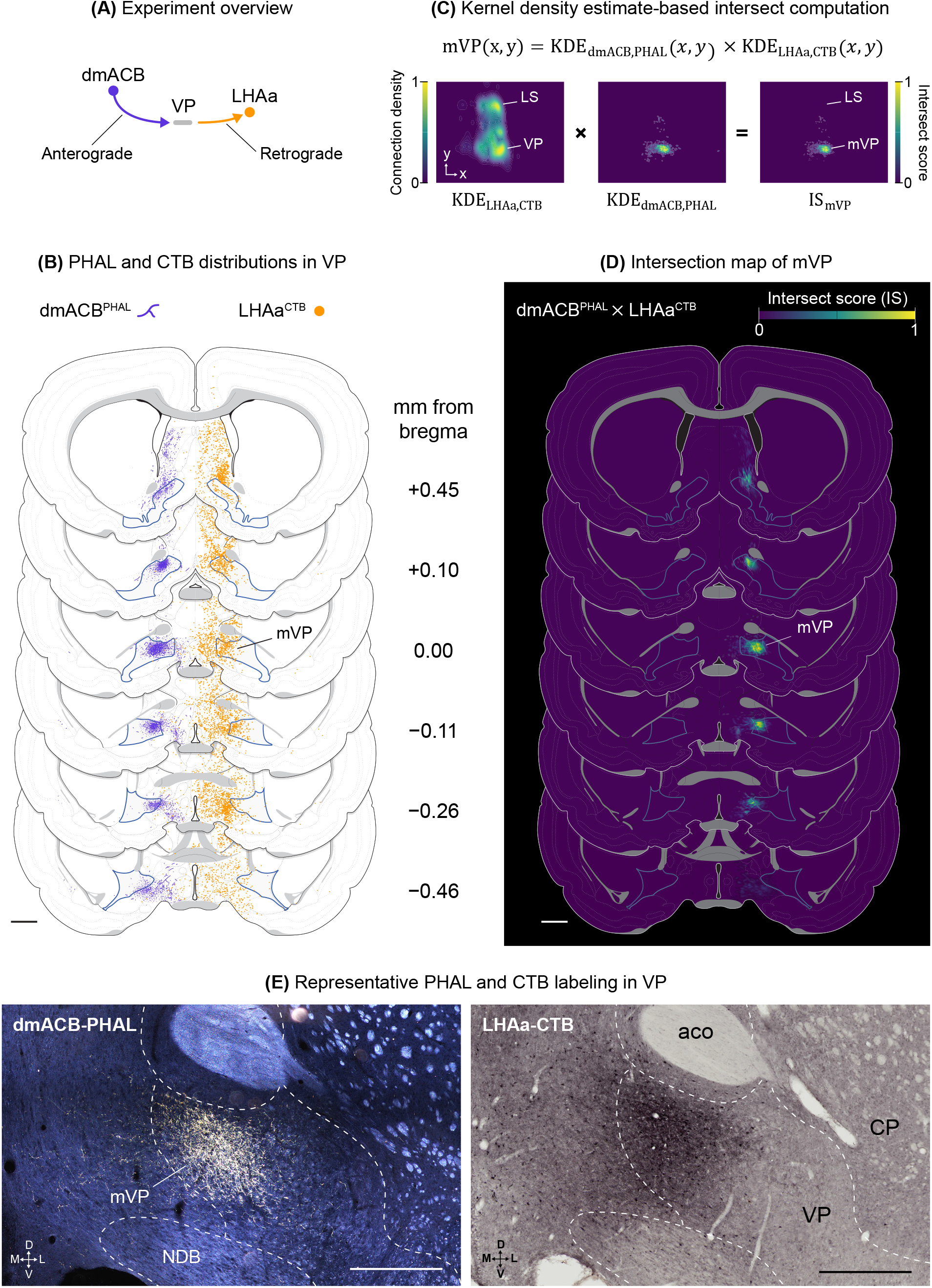
mVP as a node within a striatohypothalamic indirect pathway. **(A)** Diagrammatic representation of experiments. **(B)** Mapped distributions of dmACB axons (purple) and LHAa-projecting neurons (orange) in mVP. VP is shaded in gray and is bounded with blue outlines. Experiments are shown in opposite hemispheres for clarity. **(C)** Equation expressing the KDE product used to define the VPm and example KDEs computed for LHAa^CTB^ and dmACB^PHAL^ in an area containing the ventral pallidum. The right panel shows the normalized element-wise product of both KDEs (intersect score, IS). Note that large densities of LHAa-CTB density in LS are effectively filtered out in this process. **(D)** Atlas representations showing intersection scores derived using the equation in **(C)**. Blue lines indicate the VP boundary. Values on the left indicate AP distance from Bregma in mm for **(B)** and **(D)**. **(E)** Darkfield and brightfield photomicrographs showing VP following injections of PHAL into dmACB (left) and CTB into LHAa (right).See **Table 1** for abbreviations. Scale bars: 500 µm **(E)**, 1 mm in atlas scale **(B, D)**.

We generated intersection maps by computing grid-wise kernel density estimations (KDEs) and fitting their KDE product (intersection score) into atlas space **(Fig. 4C)**. Using equation (2), we defined the mVP as the intersection of dmACB^PHAL^ and LHAa^CTB^. The resulting maps showed a discrete cluster situated in the anteromedial end of VP **(Fig. 4D, E)**, supporting the interpretation of topographical segregation in the dmACB circuit.

Having identified the mVP component of the circuit, we next applied the same framework to define the dmACB itself. We defined the dmACB as the intersection of mVP^CTB^, LHAa^CTB^, and aPVT^PHAL^ labeling (**Fig. 5A**; eq. (3), see Materials and Methods section for boundary construction). We quantified the proportion of each region’s striatal connection density contained within the dmACB boundary. The analysis showed that the dmACB boundary accounted for a large proportion of mVP^CTB^, LHAa^CTB^, and aPVT^PHAL^ connection density, but little cVP^CTB^ connection density **(Fig. 5B)**, consistent with connectional segregation of the dmACB circuit. The resulting intersection map revealed that 95% of the intersection density was contained within a continuous column extending along the anterior-posterior axis in the dorsomedial ACB **(Fig. 5C, D)**.

**Figure 5.**
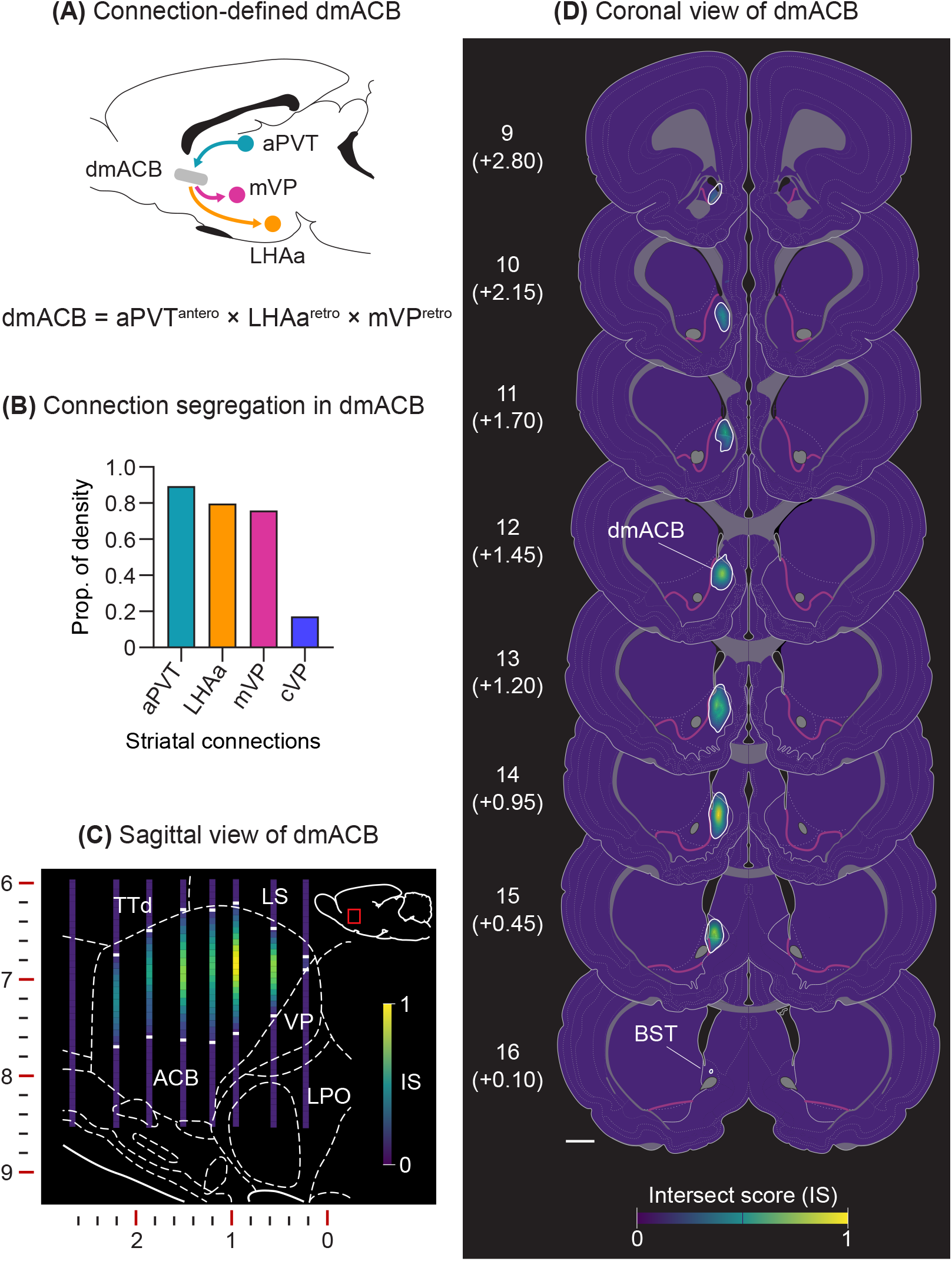
Segregation within a connection-defined dmACB. **(A)** Diagram showing neural pathways and equation expressing the KDE product used to define the dmACB. **(B)** Proportion of total striatal connection density (KDE) contained within the dmACB boundary. KDE distributions were generated using CTB labeling from LHAa, mVP, and cVP, and PHAL labeling from aPVT. **(C)** Sagittal view of the normalized intersect score (IS) for dmACB at 0.9 mm from midline. **(D)** Coronal atlas maps showing the normalized IS for dmACB. White outlines in **(C)** and **(D)** show the 95% credible region for dmACB. Solid purple lines indicate the calbindin-based ACB core-shell boundaries (see Fig. 1). Scale bars: 1 mm in atlas scale **(D)**. See **Table 1** for abbreviations.

Together, these analyses define a topographically segregated dmACB column linking aPVT inputs with outputs to mVP and LHAa. They also identify the corresponding mVP subregion that receives dmACB projections and relays information to LHAa, providing a connectional framework for the dmACB circuit.

#### Anterior-posterior differences in afferent inputs to dmACB

Our findings so far describe a connectionally uniform dmACB column. This appears inconsistent with previously reported anteroposterior differences following pharmacological manipulation of the dmACB (Reynolds & Berridge, 2003). To address this issue, we tested whether projections to the dmACB vary along its anteroposterior axis.

We analyzed non-overlapping CTB injections at the anterior and posterior extremes of the dmACB **(Fig. 6A)**. Injections targeting the anterior (a-dmACB) and posterior (p-dmACB) dmACB were comparable in size and restricted to the connection-defined boundaries of the dmACB **(Fig. 6B)**. A complete distribution of retrogradely labeled neurons for both a-dmACB and p-dmACB cases is provided in **Fig. 6C**.

**Figure 6.**
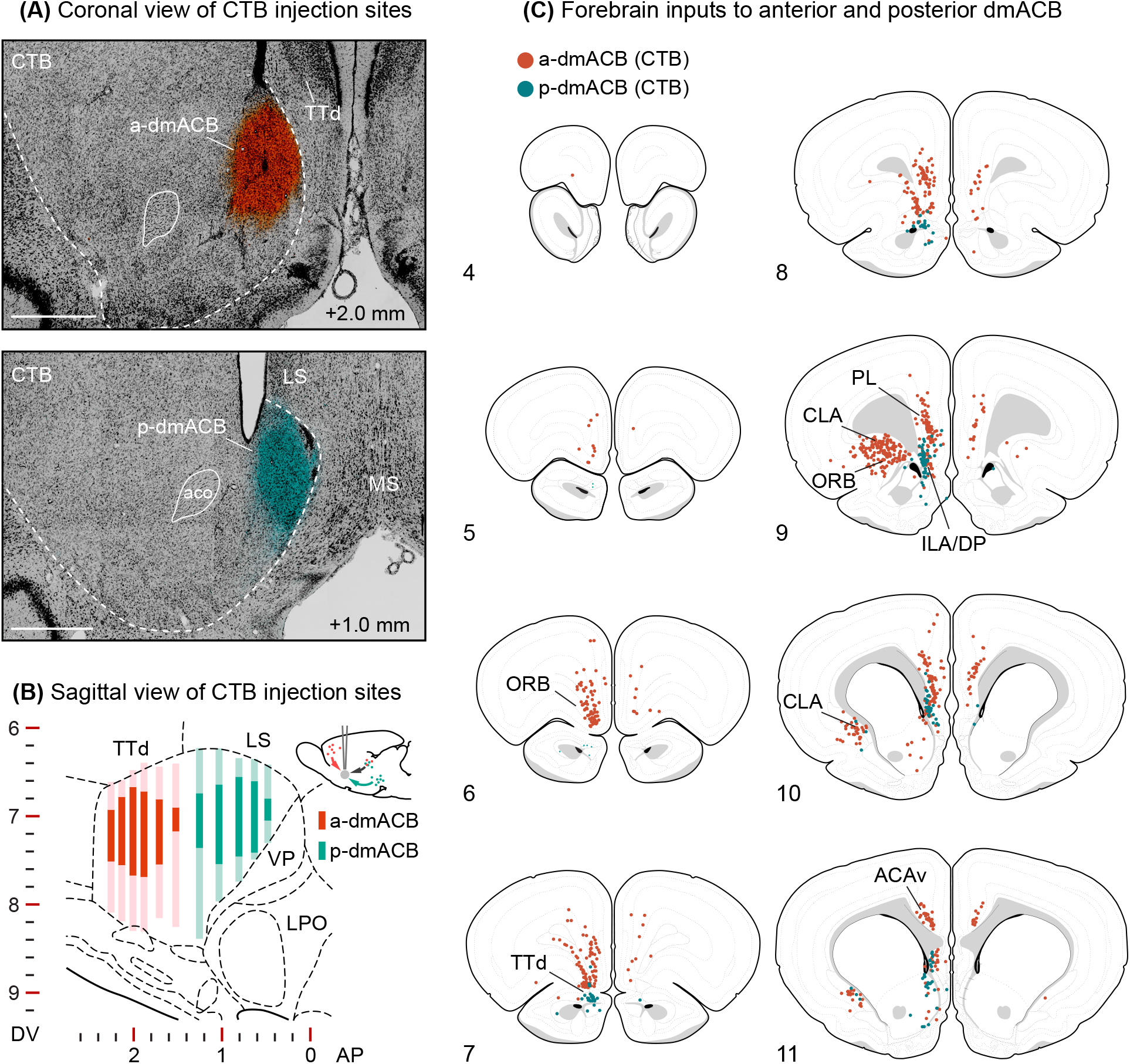
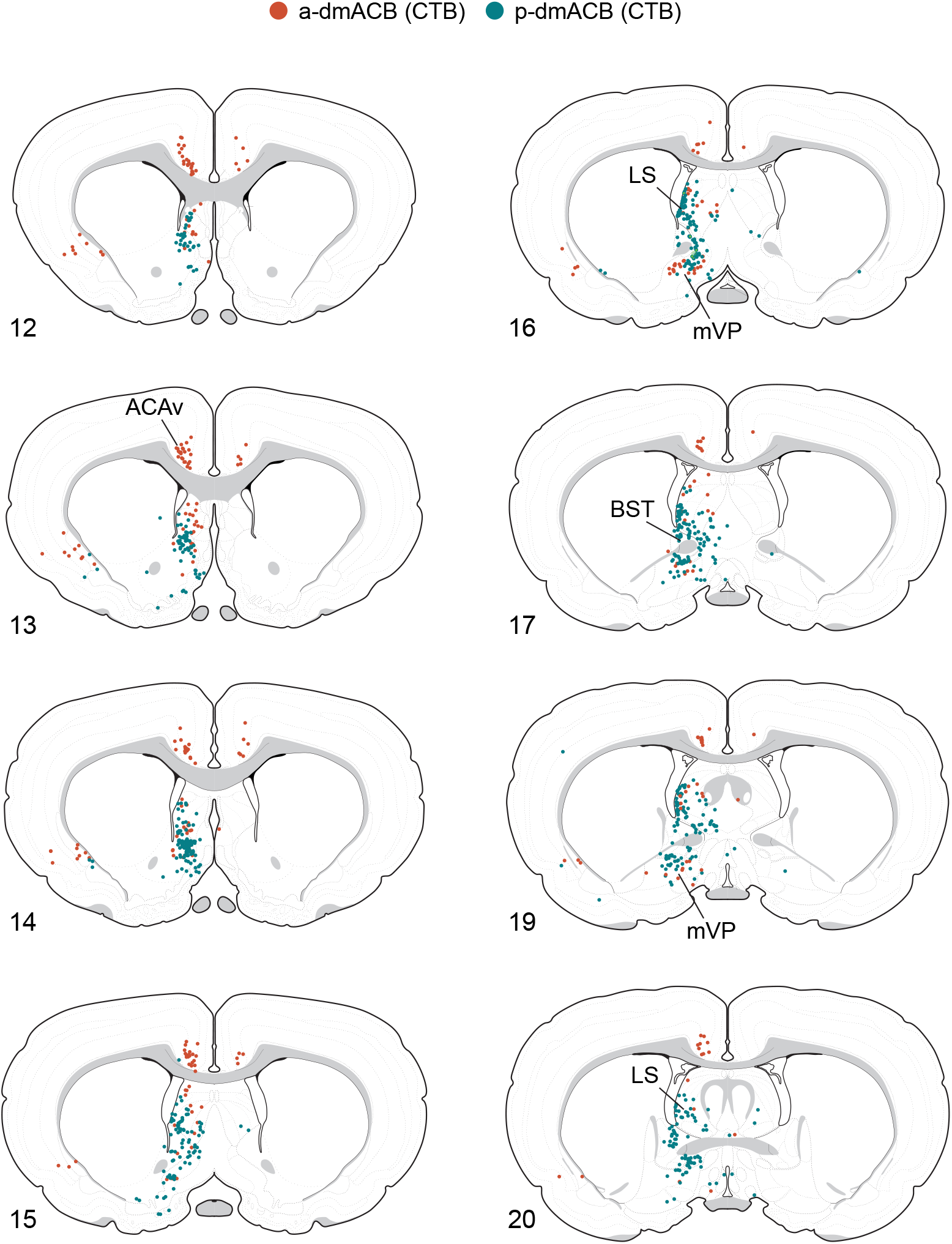
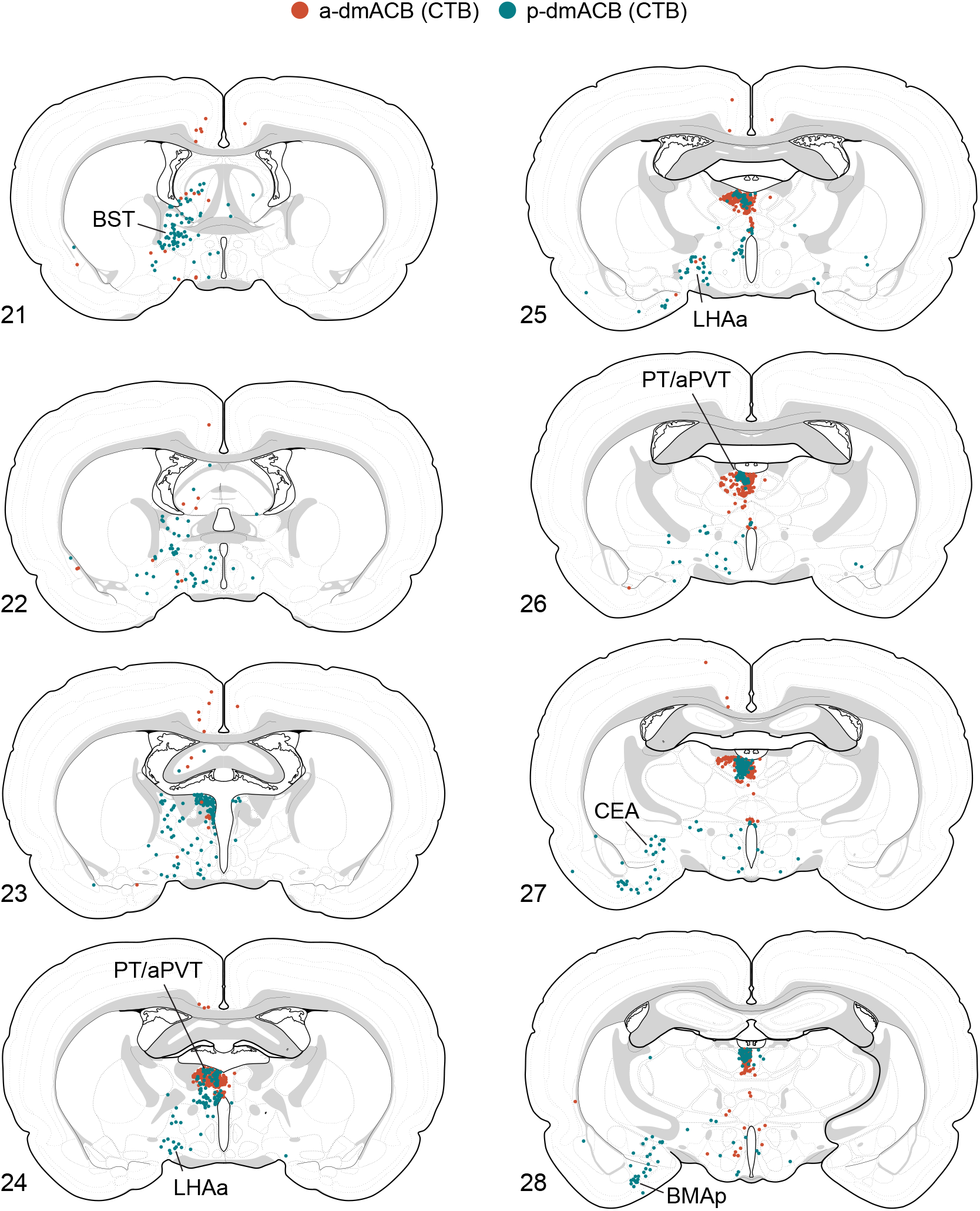
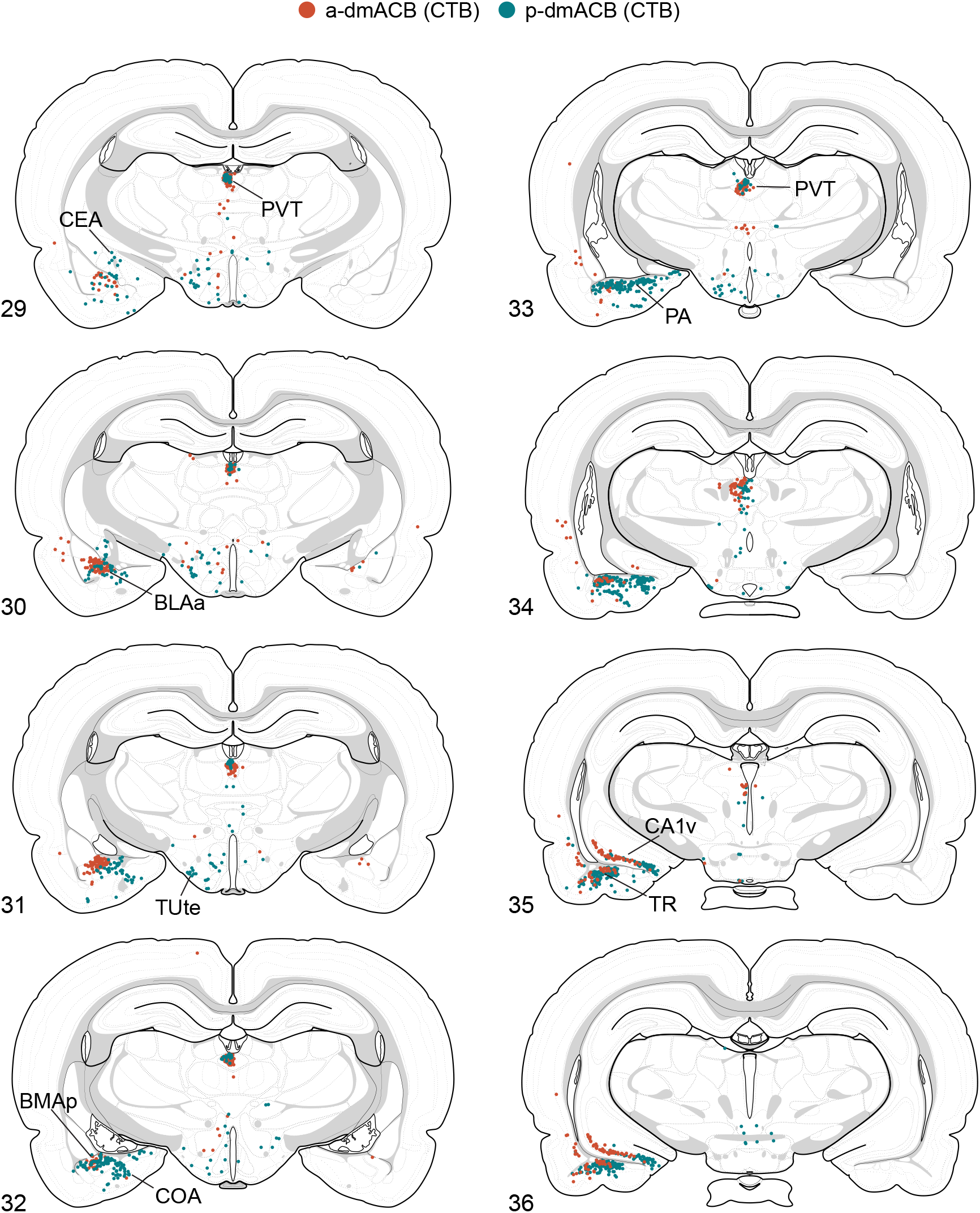
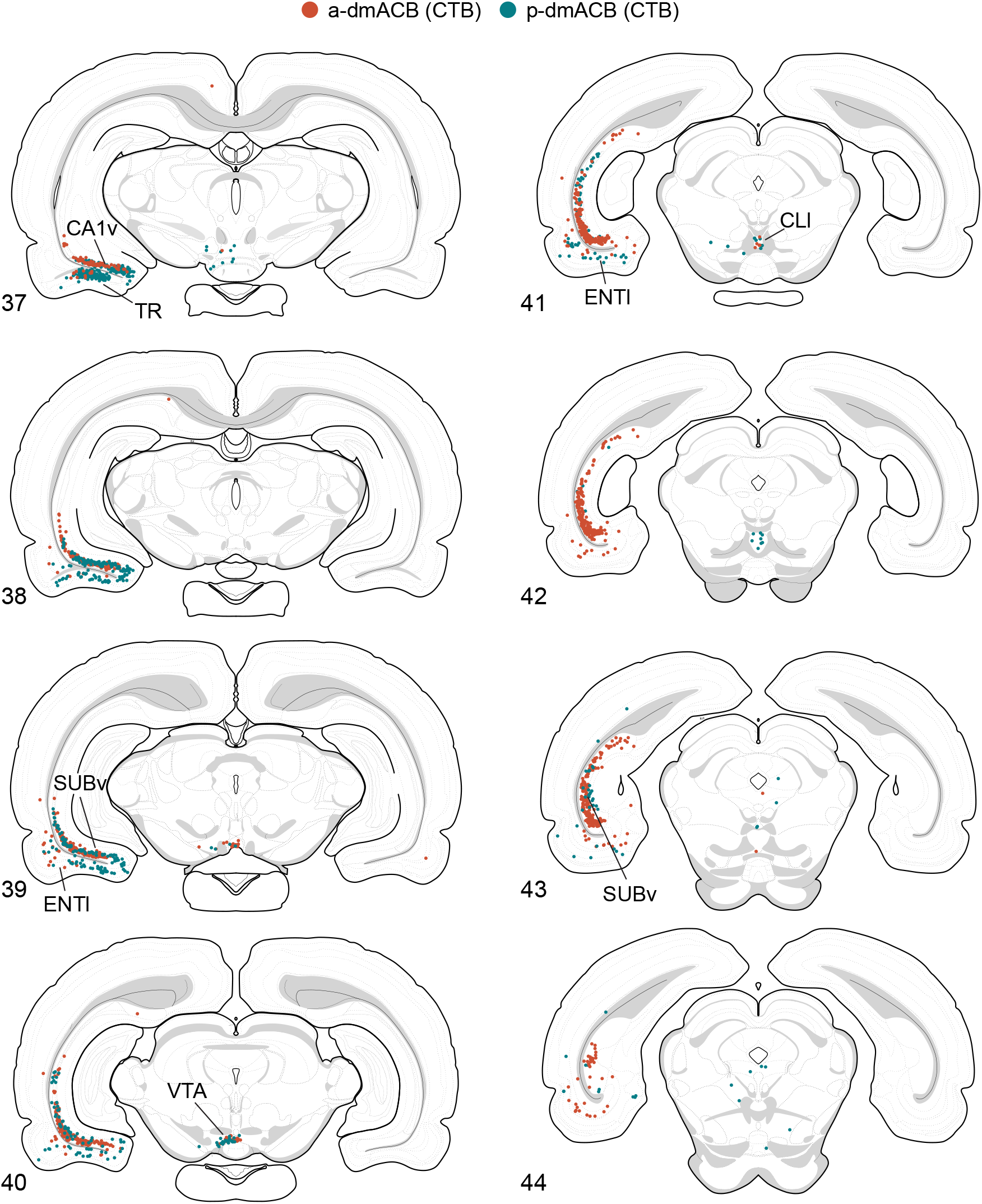
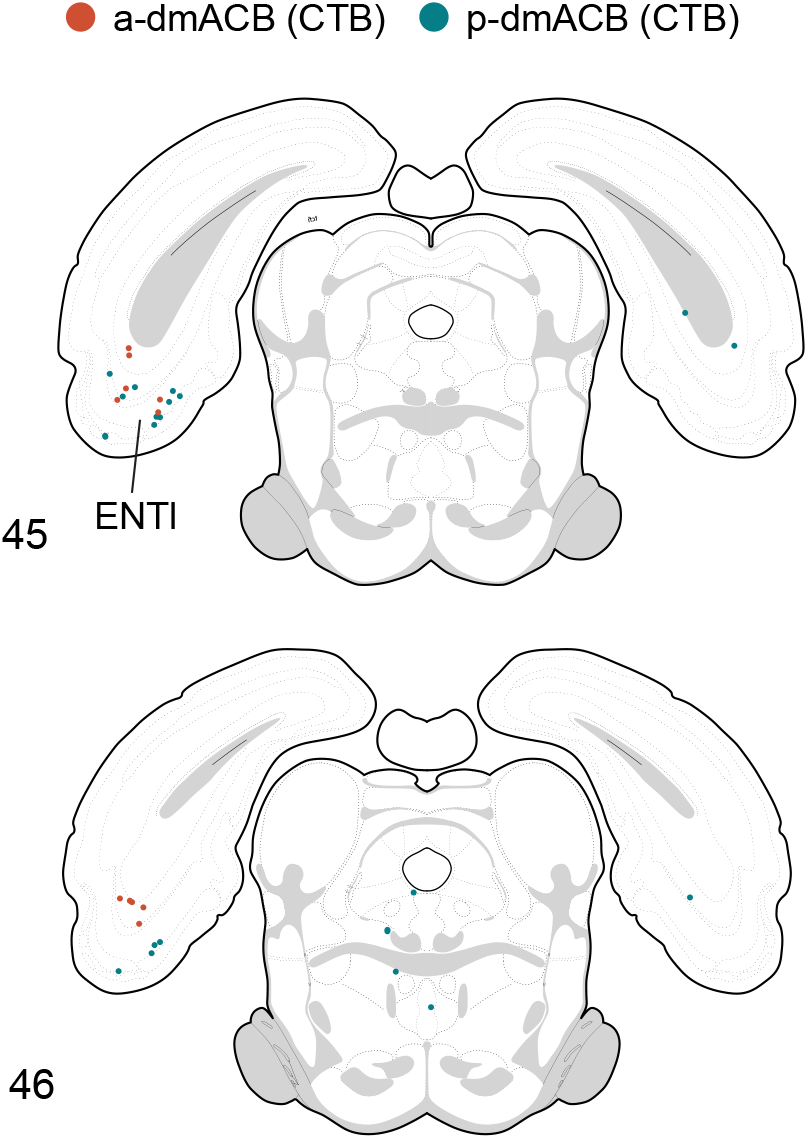
Forebrain-wide retrograde tracing of anterior and posterior dmACB. **(A)** Photomicrographs showing pseudo-colored CTB injection sites and Neurotrace staining in (above) anterior (a-dmACB) and (below) posterior (p-dmACB) dmACB. Values indicate AP distance from bregma. **(B)** Sagittal view of CTB spread after injections into a-dmACB and p-dmACB. Axes in **(B)** indicate mm distance from bregma at 0.9 mm from midline. **(C)** Mapped distributions of CTB-labeled neurons following injections into a-dmACB (red dots) and p-dmACB (green dots). Numbers in the lower-left indicate atlas levels from *Brain* Maps 4*.0*. The representative histology is shown in Fig. 7. Scale bar: 1 mm **(A)**. See **Table 1** for abbreviations.

Below, we highlight both similarities and differences in connectivity across anterior and posterior levels of the dmACB. The results are presented in an anterior-to-posterior sequence through the brain. For each immunolabeled image, the corresponding Nissl-stained section is shown in **Fig. 7**. Section headings identify key regions of interest and serve as general anatomical reference points for neighboring structures visible in the same coronal sections.

**Figure 7.**
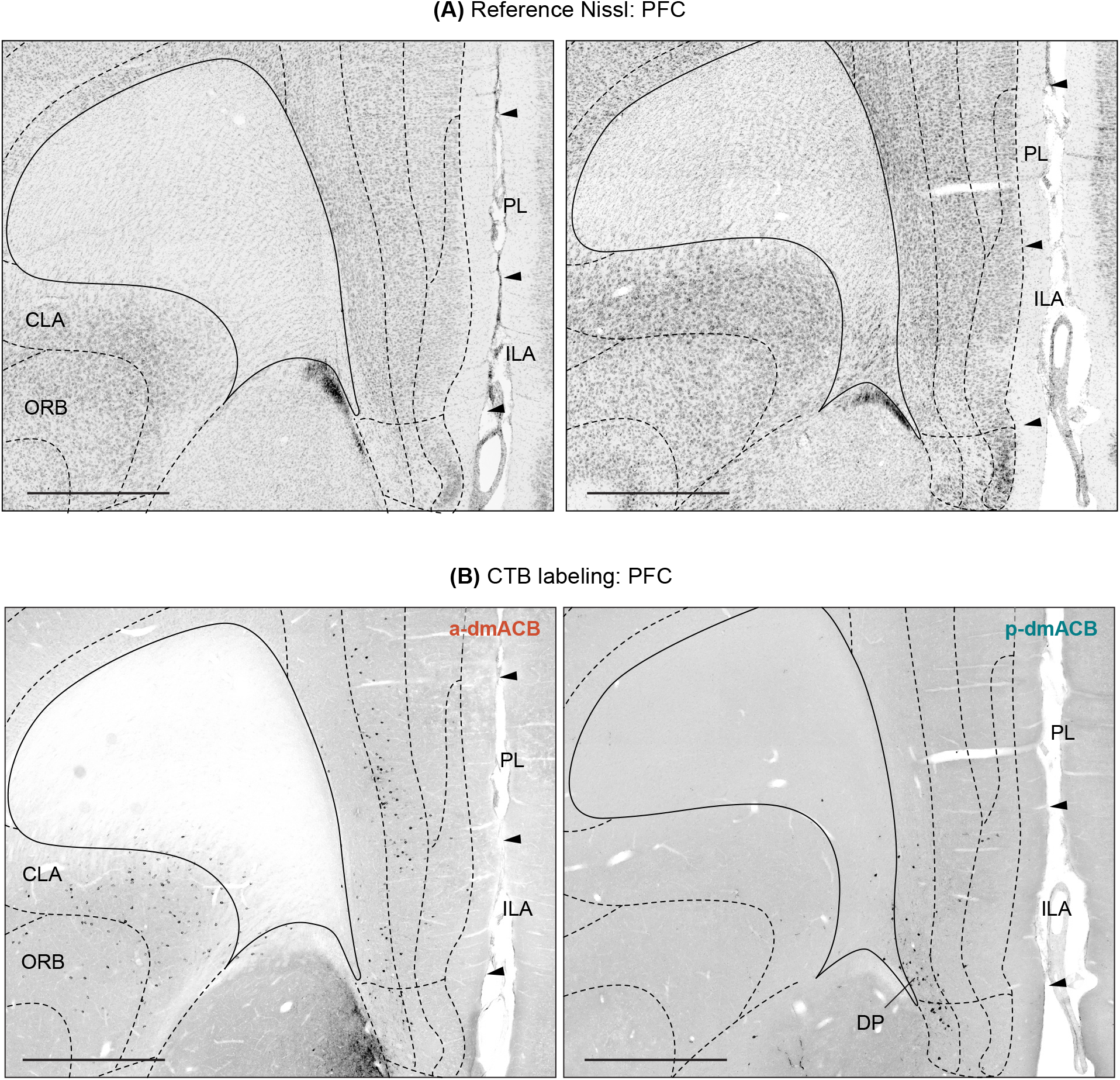
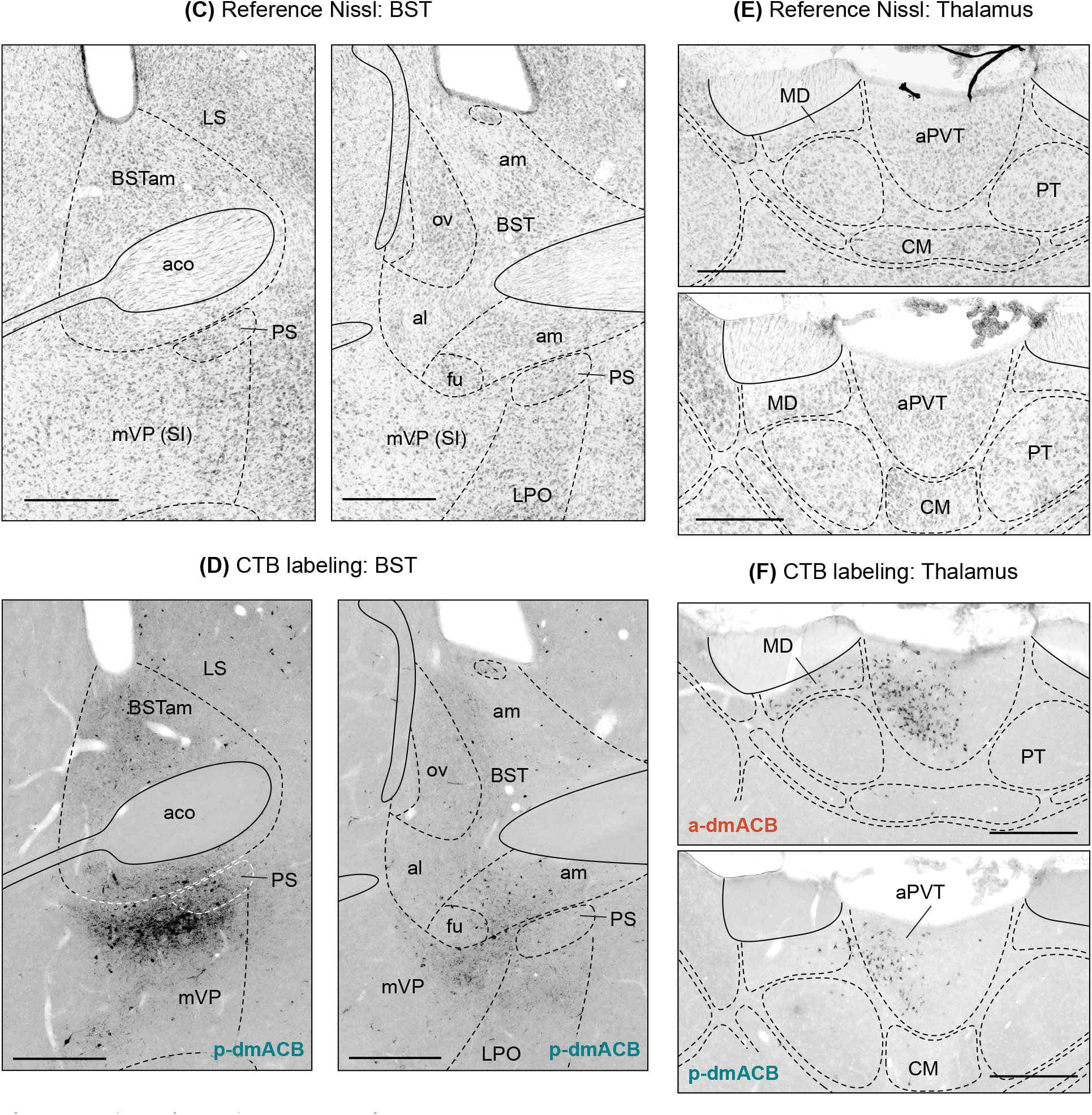
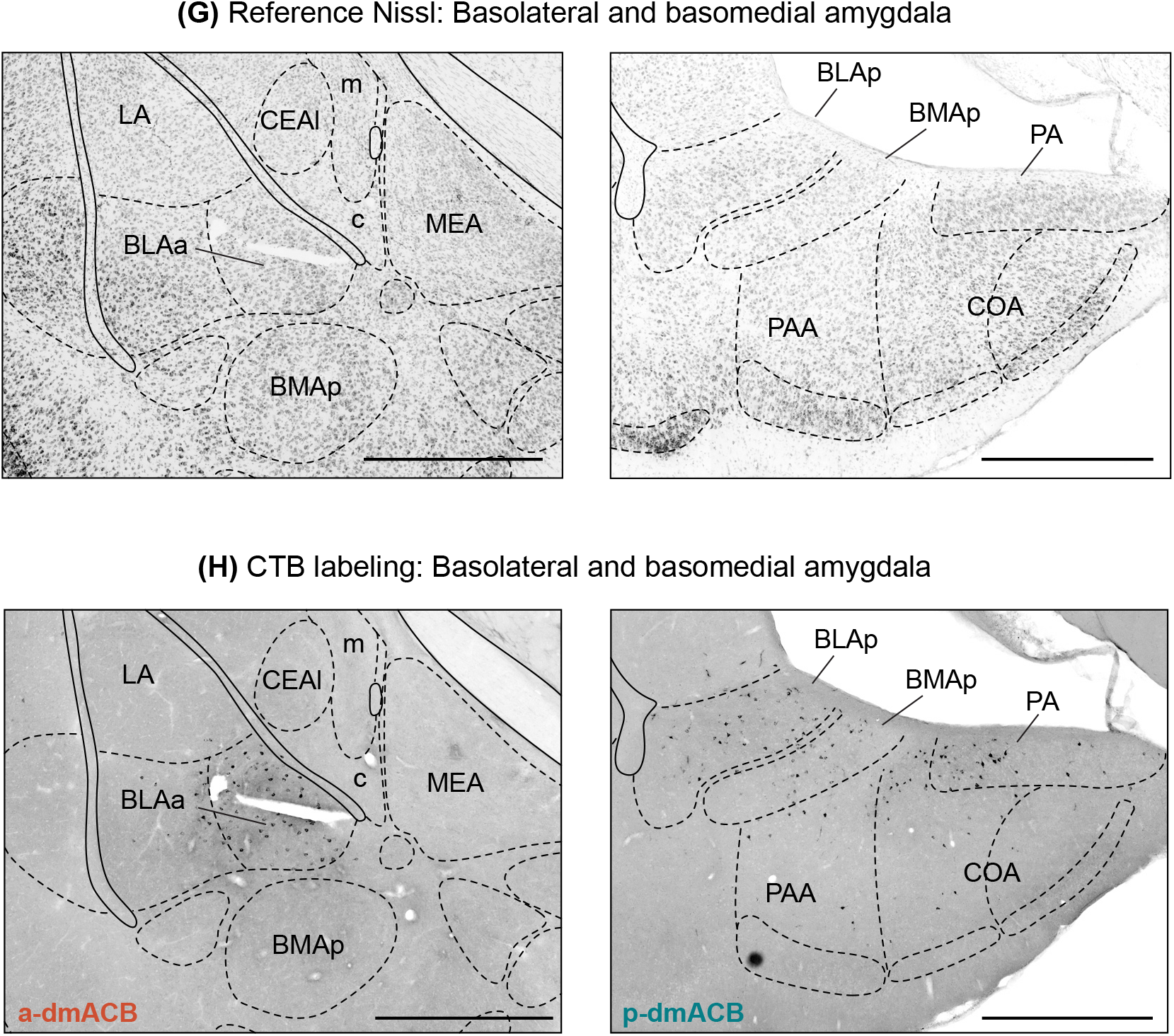
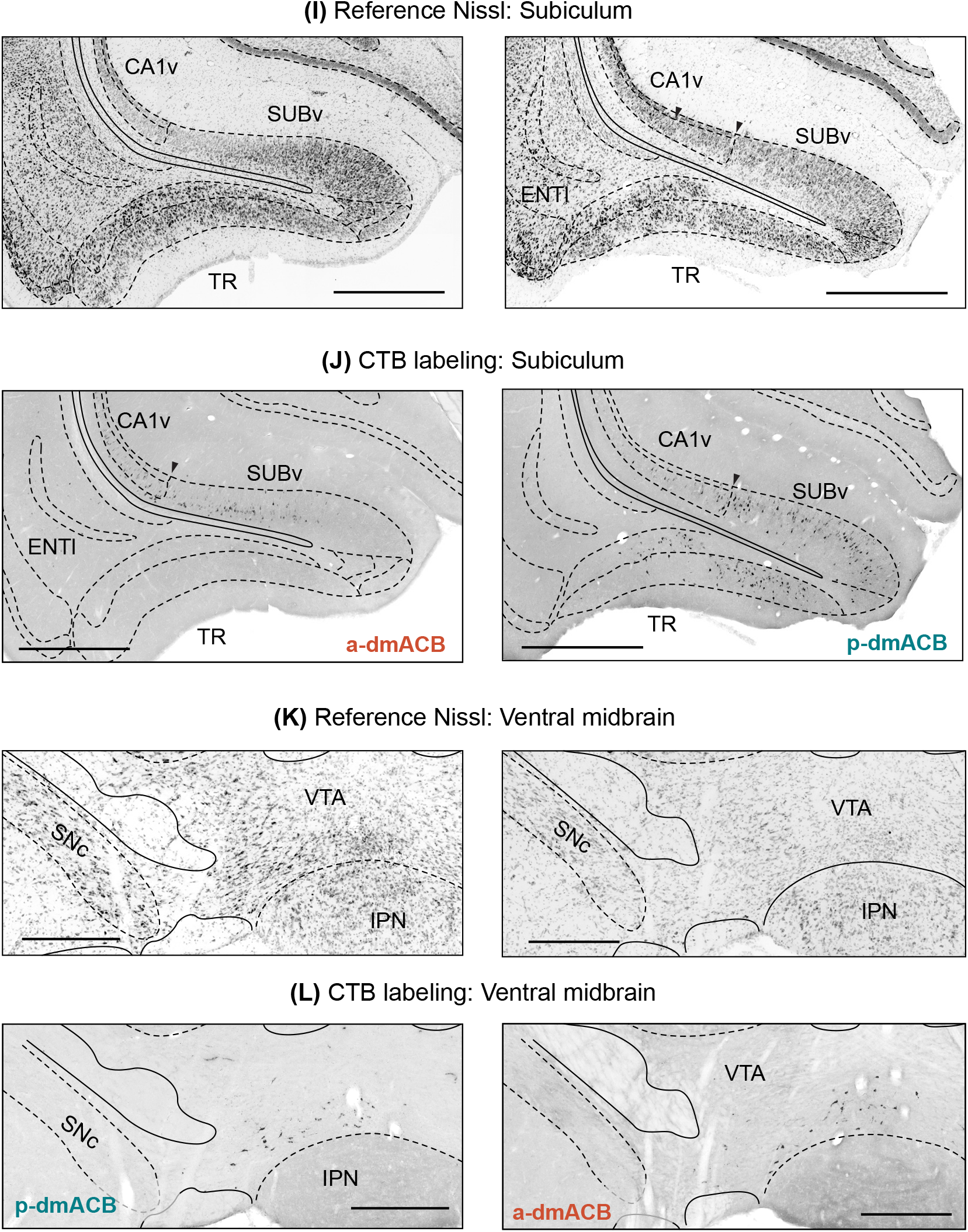
Reference Nissl parcellations and CTB labeling. Photographs showing brain region parcellations and CTB labeling used to construct maps shown in Figure 6C. Boundaries were identified using definitions and nomenclature from *Brain* Maps 4*.0* applied to pseudo-Nissl stained sections labeled with NeuroTrace. The paired CTB image is shown with the boundaries overlaid. Representative Nissl-CTB image pairs are shown for (*above*: **(A, B)** prefrontal cortex; *next page:* **(C, D)** BST, and **(E, F)** dorsal thalamus; *page 33:* **(G, H)** amygdala; *page 34:* **(I, J)** subiculum, and **(K, L)** ventral midbrain). Scale bars: 500 µm **(C, D, E, F, K, L)**, 1 mm **(A, B, G, H, I, J)**. See **Table 1** for abbreviations.

**Figure 8.**
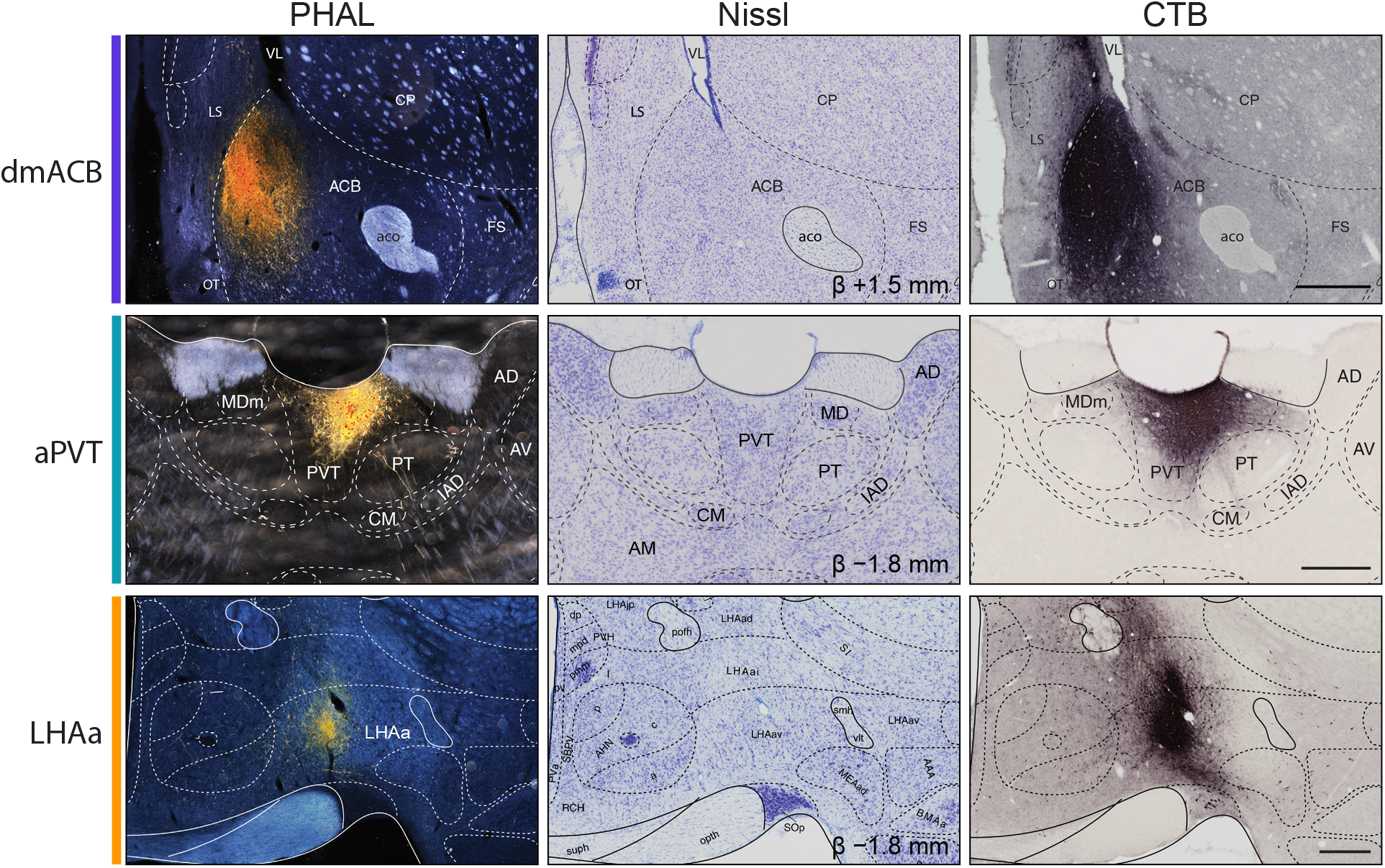
Histological localization of co-injected anterograde and retrograde tracers. Photomicrographs showing immunodetected CTB (*left column*), PHAL (*right column*), and their corresponding Nissl-stained sections (*middle column*). (*Top row*) Representative injection site in dmACB at approximately +1.5 mm from Bregma. (*Middle row*) Representative injection site in aPVT at approximately −1.8 mm from Bregma. (*Bottom row*) Representative injection site in LHAa at approximately −1.8 mm from Bregma. Superimposed boundaries are based on the cytoarchitecture shown in the Nissl-stained sections. Scale bars: 500 µm.

### Frontal cortex

We found that our observed CTB labeling in frontal cortex predominantly arose from a-dmACB **(Fig. 6C)**. Dense CTB labeling from was observed in *ventral* (ORBv) and *medial* (ORBm) *orbital areas (Krettek & Price, 1977)* of the *orbital region (Rolando, 1819)*, and the *prelimbic area (Brodmann, 1909)* (PL) **(Fig. 7A, B)**. This region also formed a moderate density of connections with the *anterior cingulate area ventral part (Krettek & Price, 1977)* (ACAv) and *infralimbic area (Rose & Woolsey, 1948)* (ILA).

The a-dmACB received some inputs from *claustrum (Burdach, 1822)* (CLA) and *agranular insular area (Swanson, 2018)* (AI). In contrast, p-dmACB connections were absent in these areas. Instead, we observed strong CTB labeling in *tenia tecta dorsal part (Swanson, 1992)* (TTd) and a ventral part of ILA that corresponds to the dorsal peduncular (DP) cortex **(Fig. 7A, B)**. The DP and CLA connections were common to both the a-dmACB and p-dmACB.

### Bed nuclei of terminal stria (Gurdjian, 1925) (BST)

At the level of BST, most CTB labeling arose from p-dmACB **(Fig. 6C)**. A moderate density of CTB labeling from p-dmACB was present throughout all levels of BST **(Fig. 7C, D)**. Dorsally, the *ventrolateral zone (Risold & Swanson, 1997)* (LSr.vl) of the *lateral septal nucleus (Ramón y Cajal, 1909–1911)* (LS) was also labeled mostly with CTB from p-dmACB. Consistent with our dmACB framework, CTB labeling was specifically found in mVP after injections into a-dmACB and p-dmACB.

### Basolateral amygdalar complex (Swanson, 2015) (BLX)

As with the BST, we found that labeling in the BLX came from p-dmACB **(Fig. 6C)**. Low to moderate densities of CTB labeling from p-dmACB was present throughout the structure. Within the BLX, most CTB labeling from the p-dmACB was detected in the *posterior amygdalar nucleus (Canteras et al., 1992)* (PA) and in the posterior parts of *basomedial* (BMAp) and *basolateral amygdalar nuclei (>1840)* (BLAp) **(Fig. 7G, H)**. A moderate density of p-dmACB CTB labeling was detected in *cortical amygdalar area (>1840)* (COA) and the *medial part* of the *central amygdalar nucleus (McDonald, 1982)* (CEAm). Although a-dmACB CTB injections did not widely label the amygdala, we observed a dense cluster in the BLAp **(Fig. 7G, H)**. The cluster had some spatial overlap with CTB labeling from p-dmACB, but p-dmACB appeared to label a topographically distinct space in the posterior end of BLAp.

In diencephalon, CTB from the a-dmACB and p-dmACB densely labeled the anterior-posterior extent of PVT **(Fig. 7E, F)**. This finding agrees with our connectional definition of dmACB and further suggests that projections from posterior PVT can be included as defining criteria. The adjacent *paratenial nucleus (>1840)* (PT) was densely labeled with CTB from a-dmACB but not p-dmACB **(Fig. 6C)**. Low to moderate densities of CTB labeling from p-dmACB and a-dmACB were present in several hypothalamic nuclei. CTB from p-dmACB was detected in medial aspects of the tuberal LHA, *ventral premammillary nucleus (>1840)* (PMv), and the *terete part (Petrovich et al., 2001)* (TUte) of the *tuberal nucleus (>1840)* (TU). CTB from a-dmACB was detected in dorsomedial parts of the *ventromedial hypothalamic nucleus (>1840)* (VMH).

### Hippocampal formation (Swanson & Cowan, 1977) (HF)

At the level of HF, most CTB labeling we observed came from a-dmACB projections arising from the *subiculum (>1840)* (SUB), particularly its *ventral part (Swanson & Cowan, 1975)* (SUBv) **(Fig. 6C)**. CTB injections into a-dmACB and p-dmACB identified the anterior SUBv as a likely zone of overlap and, hence, a potential defining criterion for dmACB **(Fig. 7I, J)** on hodological grounds. Inputs to a-dmACB and p-dmACB similarly overlapped in the posterior region of *Field CA1 (Lorente de Nó, 1934)* (CA1), specifically its *ventral part (Risold & Swanson, 1996)* (CA1v). CTB injections into p-dmACB showed labeling in the nearby *postpiriform transition area (Canteras & Swanson, 1992)* (TR; alternatively, the amygdalopiriform transition area), whereas CTB labeling from a-dmACB was not present **(Fig. 7I, J)**. More caudal sections showed some CTB labeling from a-dmACB and p-dmACB in the *lateral entorhinal area (>1840)* (ENTl).

Within the *midbrain (von Baer, 1837)*, the *ventral tegmental area (Tsai, 1925)* (VTA) presented as another zone of mesoscale connection overlap **(Fig. 7K, L)**. CTB labeling from a-dmACB and p-dmACB was restricted to the ventromedial part of the VTA and *interfascicular raphe nucleus (>1840)* (IF) bordering the *interpeduncular nucleus (>1840)* (IPN). Based on their similar CTB distributions, the ventromedial VTA and IF can potentially serve to help define dmACB on hodological grounds although their cell-type isolation may be a challenge. More caudally, we observed CTB labeling from p-dmACB in the *central linear raphe nucleus (>1840)* (CLI).

### Cortical components of the dmACB circuit

Cortical components of basal ganglia circuits are characterized by thalamocortical inputs and corticostriatal outputs (Alexander et al., 1986). Accordingly, we defined cortical components of the dmACB circuit as regions receiving aPVT projections while projecting to both dmACB and LHAa (eq. 4). We selected representative injections restricted to each region **(Fig. 8)** and applied these criteria throughout the cortex, identifying three major cortical convergence zones.

We first examined the medial prefrontal cortex (mPFC) for the dmACB circuit criteria **(Fig. 9A)**. Each criterion was satisfied for the deep layers of ventral mPFC **(Fig. 9B)**. CTB labeling from the dmACB and LHAa was found throughout the mPFC, including *anterior cingulate (Brodmann, 1909)* (ACA), PL, and ILA **(Fig. 9C)**. Retrograde labeling from dmACB and LHAa was mainly detected within layer 5 the anteroposterior extent of the ventral mPFC. In contrast, PHAL-labeled aPVT axons were limited to a narrow space in the caudal half of mPFC, mainly along the boundary between the PL and ILA **(Fig. 9C)**. The connectional overlap zone can be localized to a ‘subgenual’ part of the *cingulate region (Brodmann, 1909)*.

We next examined the *basolateral amygdalar complex (Swanson, 2015)* (BLX) for the dmACB circuit criteria **(Fig. 10A)**. Only amygdalar regions posterior to the *inferior horn of lateral ventricle (Bell, 1802)* (VLi) satisfied all three criteria **(Fig. 10B)**. PHAL labeling from aPVT and CTB labeling from dmACB and LHAa converged within the medial aspect of the posterior *basolateral amygdalar nucleus (>1840)* (BLAp), adjacent to the posterior *basomedial amygdalar nucleus (>1840)* (BMAp) **(Fig. 10C)**. In contrast, anterior BLAp lacked labeling from one or more circuit components. Likewise, although PHAL-labeled aPVT axons densely innervated the *central amygdalar nucleus (>1840)* (CEA), this region contained only sparse retrograde labeling from LHAa and no labeling from dmACB. Conversely, the *medial amygdalar nucleus (Johnston, 1923)* (MEA) contained prominent LHAa-projecting neurons but received weak aPVT input and no retrograde labeling from dmACB. Together, these observations restricted the amygdalar component of the dmACB circuit to the posterior BLAp/BMAp border.

We then examined the *hippocampal formation (Swanson & Cowan, 1977)* (HF) for the dmACB circuit criteria **(Fig. 11A)**. Injections into LHAa and dmACB labeled both CA1 and the SUBv, whereas PHAL-labeled aPVT axons were restricted to SUBv **(Fig. 11B)**. Triple labeling was concentrated within the anterior SUBv **(Fig. 11C)**. We also observed overlap in the *postpiriform transition area (Canteras et al., 1992)* (TR; amygdalopiriform transition area) and the deep layers of the *cortical amygdalar area (>1840)* (COA). In contrast, posterior CA1 and SUBv contained abundant retrograde labeling from dmACB but lacked detectable aPVT axons, whereas the *entorhinal area (Brodmann, 1909)* received dense aPVT projections but contained sparse retrograde labeling from both dmACB and LHAa. Together, these findings localized the hippocampal component of the dmACB circuit primarily to the anterior SUBv.

Finally, we integrated these regional observations using our KDE-based computational approach to identify cortical regions satisfying the dmACB, LHAa, and aPVT connectivity criteria throughout the cortex **(Fig. 12A)**. The resulting intersection maps recapitulated the anatomical observations, highlighting discrete convergence zones within the PL/ILA border, DP/TTd, the BLAp/BMAp border, and the anterior SUBv **(Fig. 12B–D)**.

### Experiment 2: dmACB neuronal activation during incubation of oxycodone seeking

#### Overview

The experiment consisted of three phases: oxycodone self-administration (14 days), electric barrier–induced abstinence (11 days), and 2-h tests of relapse to oxycodone seeking conducted before (abstinence day 1) and after barrier exposure (abstinence day 15) **(Fig. 13A)**. After perfusions and IHC staining, we examined whether incubation of oxycodone seeking was associated with time-dependent changes in neuronal activation, as detected by phospho-Ser^32^-Fos immunostaining (Cruz et al., 2013), in the dmACB and adjacent striatal regions.

#### Oxycodone self-administration

The male and female rats acquired oxycodone self-administration; over time, males self-administered more oxycodone than females **(Fig. 13B)**. Analysis of the number of infusions, with Sex (male, female) as a between-subjects factor and Session (1-14) as a within-subjects factor, showed significant effects of Sex (F_(1,26)_ = 18.1, p < 0.001), Session (F_(3.6,94.1)_ = 26.4, p < 0.001), and a Session × Sex interaction (F_(3.6,94.1)_ = 5.9, p < 0.001). Analysis of active lever presses, with Sex as a between-subjects factor, Session as a within-subjects factor, and inactive lever-presses as a covariate, showed significant effects of Sex (F_(1,25)_ = 8.1, p = 0.009), Session (F_(1.7,42.0)_ = 4.9, p = 0.017), and a Session × Sex interaction (F_(1.7,42.0)_ = 4.0, p = 0.031).

#### Electric barrier-induced abstinence

During the electric barrier phase, progressively increasing shock intensity suppressed oxycodone self-administration, with a stronger effect in females **(Fig. 13C)**. Analysis of infusions, with Sex as a between-subjects factor and Session (1-11) as a within-subjects factor, showed significant effects of Sex (F_(1,12)_ = 19.5, p < 0.001), Session (F_(10,120)_ = 125.3, p < 0.001), and a Session × Sex interaction (F_(10,120)_ = 15.0, p < 0.001). Analysis of active lever presses, with Sex as a between-subjects factor, Session as a within-subjects factor, and inactive lever-presses as a covariate, showed significant effects of Sex (F_(1,11)_ = 5.9, p = 0.034), Session (F_(10,110)_ = 14.3, p < 0.001), and a Session × Sex interaction (F_(10,110)_ = 4.9, p < 0.001).

#### Relapse (incubation) test

Rats tested after 15 abstinence days showed higher active lever responding than rats tested on abstinence day 1, with a somewhat stronger ‘incubation’ effect in males **(Fig. 13D)**. Analysis of total non-reinforced active lever presses, with Sex as a between-subjects factor, Abstinence Day (1, 15) as a within-subjects factor, and inactive lever responding as a covariate, showed significant effects of Sex (F_(1,23)_ = 9.9, p < 0.001) and Abstinence Day (F_(1,23)_ = 32.0, p < 0.001), but no significant Sex × Abstinence Day interaction (F_(1,23)_ = 3.4, p = 0.079). Analysis of the time course of active lever presses during the relapse tests, which included Session Time (30, 60, 90, 120 min) as an additional within-subjects factor, showed significant effects of Session Time (F_(1.3,30.6)_ = 34.2, p < 0.001), Abstinence Day × Session Time (F_(1.3,30.6)_ = 11.7, p < 0.001), and Sex × Session Time (F_(1.3,30.6)_ = 6.2, p = 0.012).

Together, these behavioral results replicate our previous findings on incubation of oxycodone seeking after electric barrier-induced abstinence in both sexes (Fredriksson et al., 2020; Fredriksson et al., 2023; Negishi et al., 2025). However, in contrast to our prior studies, **Experiment 2** showed higher oxycodone self-administration, slower suppression by the electric barrier, and somewhat greater incubation in males than in females. The reasons for these sex differences are unknown.

#### Fos immunohistochemistry

We counted Fos-labeled cells in three groups: (1) rats tested on abstinence day 1, with brains collected immediately after the 2-h test (n=9); (2) rats tested on abstinence day 15, with brains collected immediately after the 2-h test (n=9); and (3) No-Test rats, with brains collected on abstinence day 2 (n=5) or day 15 (n=5; total n=10). Because counts of Fos+ cells did not differ between the two No-Test subgroups, we combined them for statistical analyses. We initially detected Fos+ nuclei using a Cellpose-SAM model trained to ignore dimly labeled cells **(Fig. 13E)**. To evaluate whether segmentation introduced group-dependent biases, we generated histograms of segmented objects based on their average nuclear signal intensity. The proportion of nuclei across intensity bins was similar between groups, and signal-to-noise ratios did not differ **(Fig. 13F)**. Thus, differences in Fos+ cell counts do not reflect systematic differences in histology, image processing, or segmentation.

We used complementary approaches to define striatal regions of interest (ROIs) at AP +1.0 mm and +2.0 mm from bregma. We used calbindin immunolabeling to distinguish ACB core from shell (Jongen-Rêlo et al., 1994). We further subdivided the shell into dmACB, ventrolateral (vlACB), and ventromedial (vmACB) regions based on CTB distributions from **Experiment 1 (Fig. 13G)**. We divided the dorsal striatum (DS) into dorsomedial (DM), dorsolateral (DL), ventromedial (VM), and ventrolateral (VL) zones using geometric criteria (Morigaki et al., 2020).

Relapse testing increased Fos+ cell counts in all striatal regions in both day 1 and day 15 groups relative to the No-Test group, with the largest increases in ACB core and dmACB **(Fig. 14A, B)**. In addition, Fos+ cell counts were lower on day 15 than on day 1 in anterior and posterior dmACB, posterior vlACB, and anterior and posterior ventrolateral dorsal striatum **(Fig. 14B, C)**. We first analyzed the numbers of Fos-expressing cells using Test Condition (No-Test, Day 1 Test, Day 15 Test) as a between-subjects factor and bregma Level (anterior, posterior) and Striatal Region (four ACB and four dorsal striatal subregions) as within-subjects factors. This analysis showed a significant three-way interaction (F_(6.4,80.2)_ = 6.4, p < 0.003), along with additional significant main

Spatial representation of normalized change in Fos density from day 1 to day 15 contrasts anterior and posterior dmACB from other striatal regions **(Fig. 14D)**. Post-hoc one-way ANOVAs followed by Tukey tests (Fig. 14B,C) confirmed significant decreases in the numbers of Fos+ cells from day 1 to day 15 in anterior and posterior dmACB (F_(2,25)_ = 29.5 and 29.8, respectively, both p < 0.001; Tukey p = 0.003 and 0.050, respectively); posterior vlACB (F_(2,25)_ = 26.4, p = 0.019); anterior and posterior ventrolateral DS (F_(2,25)_ = 18.0, p = 0.022; F_(2,25)_ = 14.2, p = 0.048); and anterior dorsolateral DS (F_(2,25)_ = 17.1, p = 0.006). Spatial representation of normalized change in Fos density from day 1 to day 15 further contrasts anterior and posterior dmACB from other striatal regions **(Fig. 14D)**.

Finally, we examined the relationship between Fos+ cell counts and relapse behavior by computing Pearson’s correlation coefficients between the counts in striatal ROIs and active lever pressing during the relapse tests (both the full 2-h session and the first 30 min, when most responses occurred). We found significant negative correlations between active lever pressing and Fos+ cell counts from combined Day 1 and Day 15 test groups in anterior (r = −0.619, p = 0.006) and posterior (r = −0.504, p = 0.033) **(Fig. 14E)** dmACB during the first 30 min, and in anterior dmACB during the 2-h session (r = −0.571, p = 0.013). Fos+ cell counts were not correlated with active lever pressing in the other subregions of ACB and DS **(Table 3)**.

**Table 3.** Pearson *r* correlations between active lever presses during the relapse tests (combining days 1 and 15) and Fos expression in the different striatal regions.

| **Area** | **Bregma** | **Test condition (30 minutes)** | **Test condition (120 minutes)** |
|---|---|---|---|
| Nucleus accumbens core (ACBcore) | Anterior | Pearson r = –0.169, p = 0.502 | Pearson r = –0.091, p = 0.720 |
|  | Posterior | Pearson r = 0.096, p = 0.705 | Pearson r = 0.147, p = 0.561 |
| Dorsomedial nucleus accumbens (dmACB) | Anterior | Pearson r = –0.619, p = **0.006** | Pearson r = –0.571, p = **0.013** |
|  | Posterior | Pearson r = –0.504, p = **0.033** | Pearson r = –0.419, p = 0.083 |
| Ventromedial nucleus accumbens (vmACB) | Anterior | Pearson r = 0.025, p = 0.921 | Pearson r = –0.007, p = 0.977 |
|  | Posterior | Pearson r = 0.215, p = 0.392 | Pearson r = 0.217, p = 0.387 |
| Ventrolateral nucleus accumbens (vlACB) | Anterior | Pearson r = 0.072, p = 0.775 | Pearson r = 0.082, p = 0.747 |
|  | Posterior | Pearson r = –0.099, p = 0.695 | Pearson r = –0.163, p = 0.517 |
| Dorsal striatum dorsomedial (DM) | Anterior | Pearson r = –0.092, p = 0.716 | Pearson r = –0.045, p = 0.859 |
|  | Posterior | Pearson r = 0.301, p = 0.225 | Pearson r = 0.417, p = 0.085 |
| Dorsal striatum dorsolateral (DL) | Anterior | Pearson r = –0.425, p = 0.078 | Pearson r = –0.411, p = 0.090 |
|  | Posterior | Pearson r = –0.299, p = 0.229 | Pearson r = –0.180, p = 0.475 |
| Dorsal striatum ventromedial (VM) | Anterior | Pearson r = 0.195, p = 0.439 | Pearson r = 0.289, p = 0.244 |
|  | Posterior | Pearson r = –0.008, p = 0.974 | Pearson r = 0.025, p = 0.921 |
| Dorsal striatum ventrolateral (VL) | Anterior | Pearson r = –0.186, p = 0.461 | Pearson r = –0.171, p = 0.497 |
|  | Posterior | Pearson r = –0.039, p = 0.876 | Pearson r = –0.029, p = 0.909 |

## Discussion

We identified a subregion of the ACB, which we term the dmACB column, that forms a longitudinal structure extending along the anteroposterior axis, defined at the mesoscale by convergent anatomical connections with mVP, LHAa, and aPVT **(Fig. 15A)**. Although the column appears relatively uniform at a coarse anatomical level, its heterogeneous internal organization becomes apparent when examining afferent inputs along its anteroposterior axis **(Fig. 15B)**. Supporting this idea, forebrain CTB labeling following injections into anterior versus posterior dmACB showed large differences in anteroposterior input patterns from cortical regions, subcortical structures (including the amygdala and BST), and other areas.

**Figure 9.**
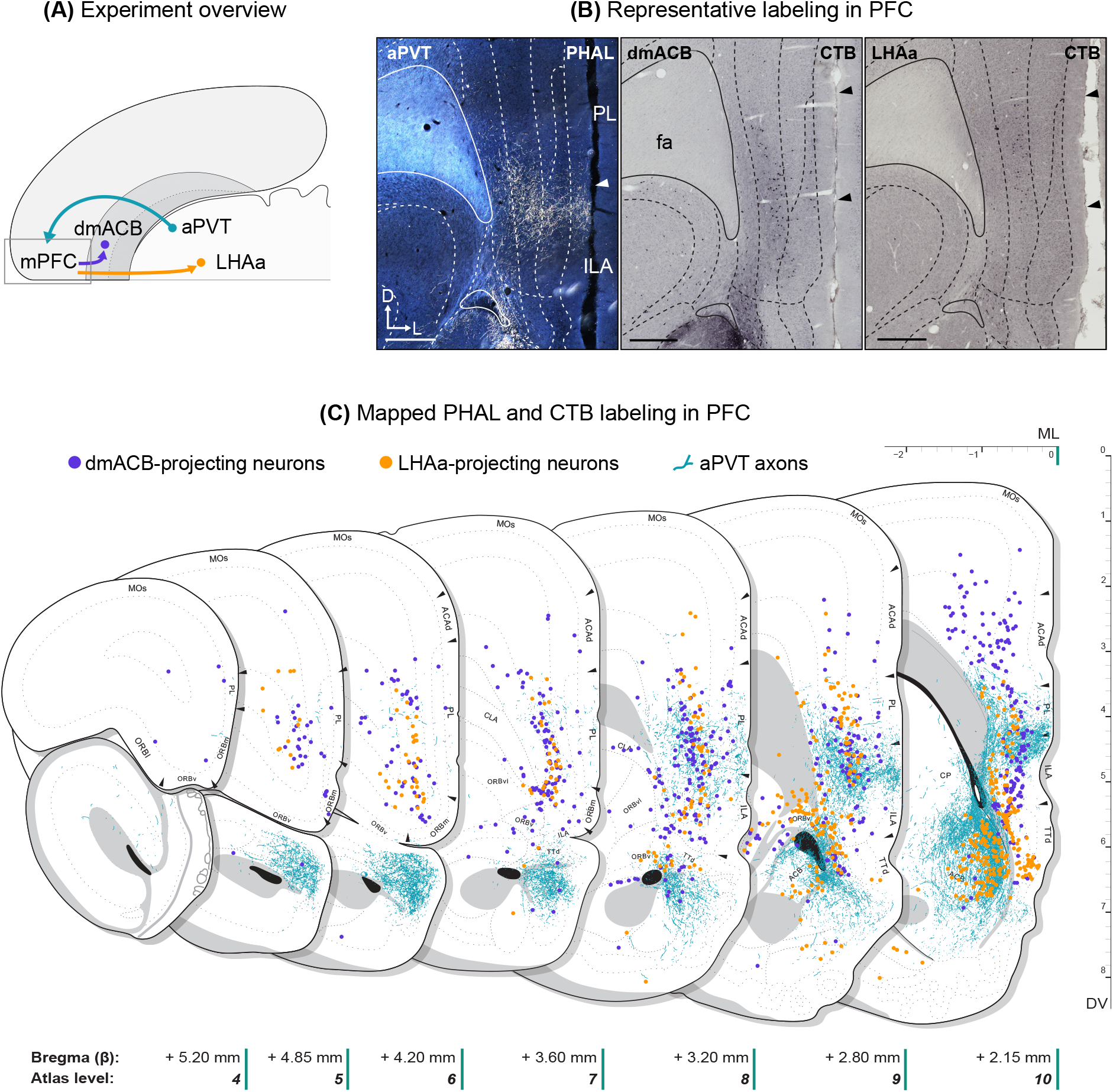
Convergent dmACB, LHAa, and aPVT connections in mPFC. **(A)** Diagrammatic representation of experiments. Circles show injected regions and arrows show the direction of the connections. **(B)** Photomicrographs show immunodetected PHAL and CTB in mPFC at approximately +3.0 mm from bregma. Injection site locations are indicated with abbreviations in the upper-left corners and tracers are indicated in the upper-right corners of each image. Arrowheads point to boundary locations between cortical areas. Boundaries were determined using adjacent Nissl-stained sections. **(C)** Mapped distributions of aPVT axons (*teal*), dmACB-(*purple*) and LHAa-projecting (*orange*) neurons in mPFC. Note the overlap of each label in a subgenual part of mPFC. Scale bar: 500 µm **(B).**

**Figure 10.**
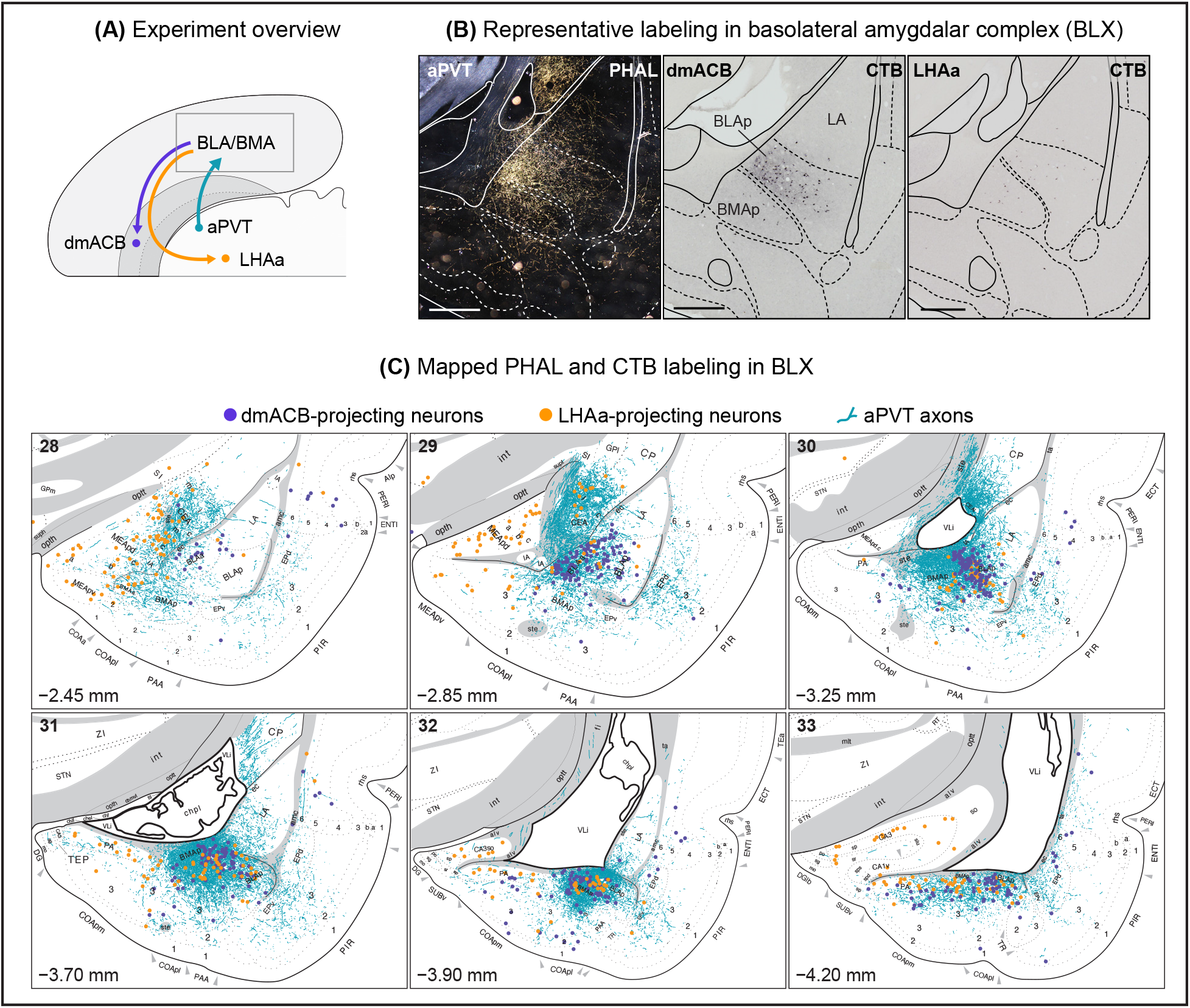
Convergent dmACB, LHAa, and aPVT connections in amygdala. **(A)** Diagrammatic representation of experiments. Circles show injected regions and arrows show the direction of the connections. **(B)** Photomicrographs showing immunodetected PHAL and CTB in amygdala at approximately −3.5 mm from Bregma. Injection site locations are indicated with abbreviations in the upper-left corners and tracers are indicated in the upper-right corners of each image. Boundaries were determined using adjacent Nissl-stained sections. **(C)** Mapped distributions of aPVT axons (*teal*), dmACB-(*purple*) and LHAa-projecting (*orange*) neurons in amygdala. Note the overlap of each label in a medial space between BLAp and BMAp. Values in lower-left corners indicate the approximate distance from Bregma and corresponding atlas levels from Swanson (Swanson, 2018) are indicated in the upper-left corners of each panel. Scale bar: 500 µm **(B)**.

**Figure 11.**
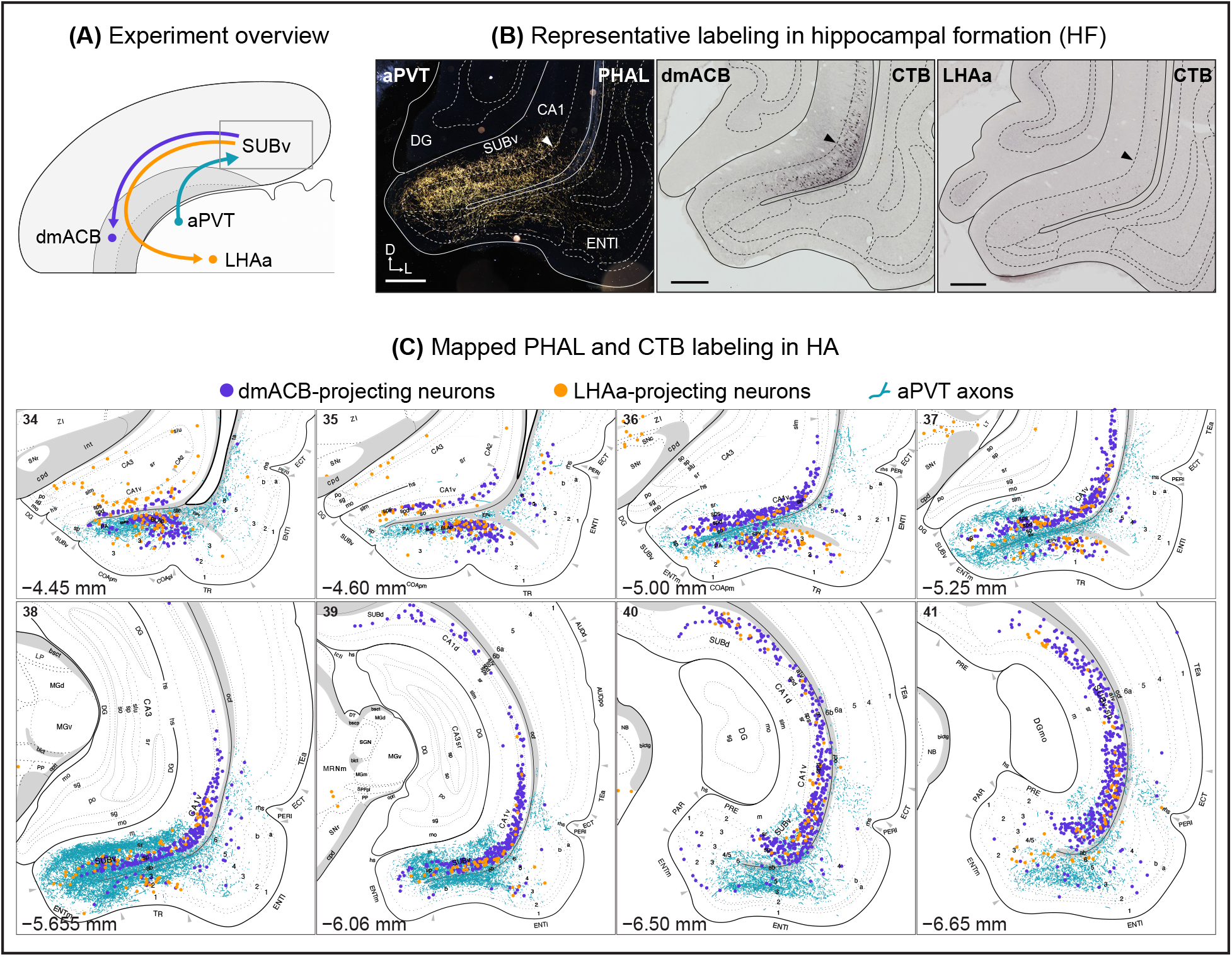
Convergent dmACB, LHAa, and aPVT connections in hippocampus. **(A)** Diagrammatic representation of experiments. Circles show injected regions and arrows show the direction of the connections. **(B)** Photomicrographs showing immunodetected PHAL and CTB in SUBv at approximately −6.0 mm from Bregma. Injection site locations are indicated with abbreviations in the upper-left corners and tracers are indicated in the upper-right corners of each image. Note the overlap of each label in a lateral aspect of the anterior SUBv. **(C)** Mapped distributions of aPVT axons (*teal*), dmACB-(*purple*) and LHAa-projecting (*orange*) neurons in hippocampus. Values in lower-left corners indicate the approximate distance from Bregma and corresponding atlas levels from Swanson (Swanson, 2018) are indicated in the upper-left corners of each panel. See Abbreviations for nomenclature. Scale bar: 500 µm **(B).**

**Figure 12.**
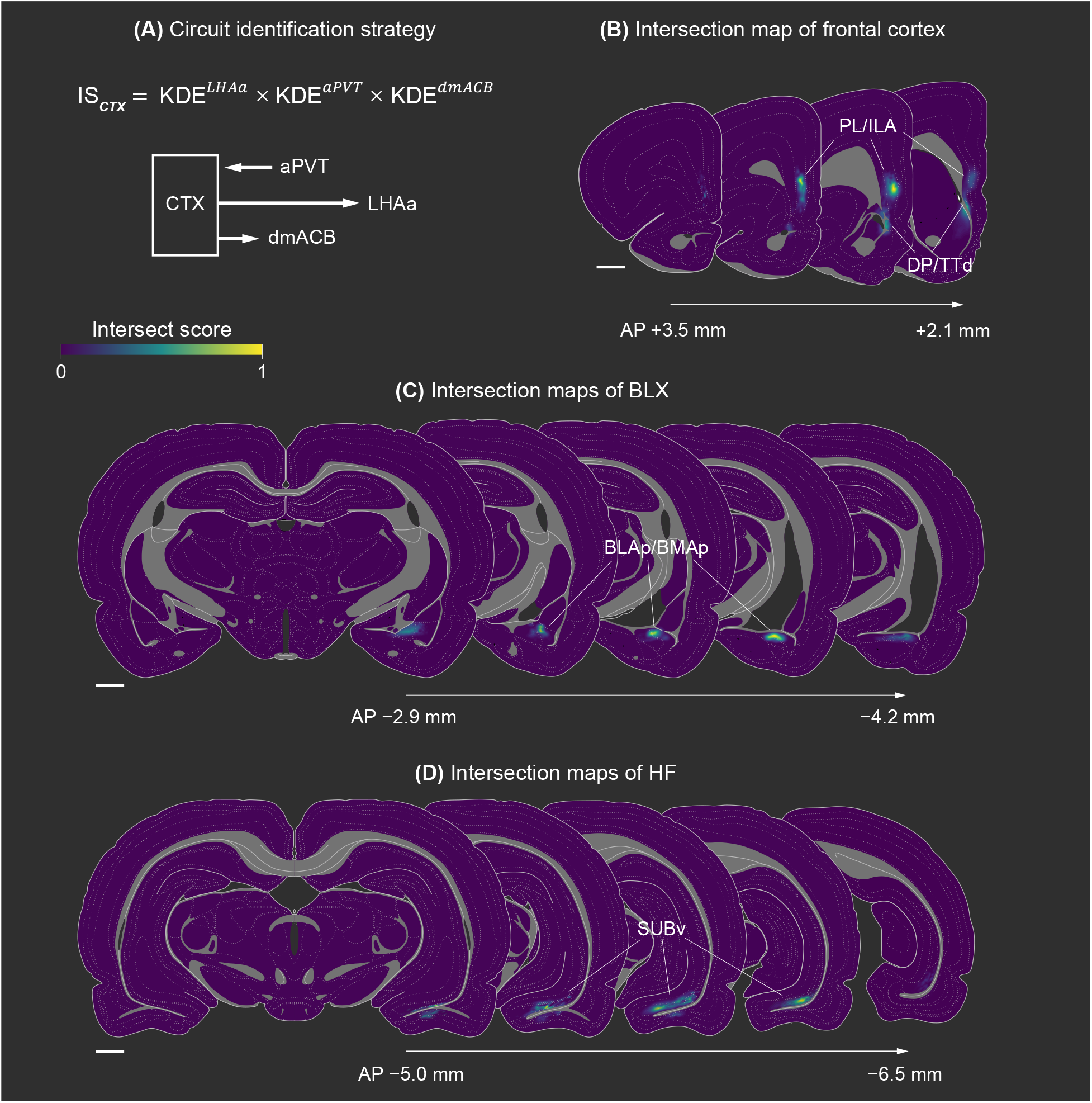
Cortical regions of the dmACB circuit. **(A)** Equation and schematic expressing the KDE product and neural pathways used to identify cortical regions (CTX). Intersection maps computed using the equation in **(A)** are shown for **(B)** frontal cortex, **(C)** basolateral amygdalar complex, and **(D)** hippocampal formation. AP values denote distance from bregma. Scale bar: 1 mm in atlas scale **(B, C, D).**

**Figure 13.**
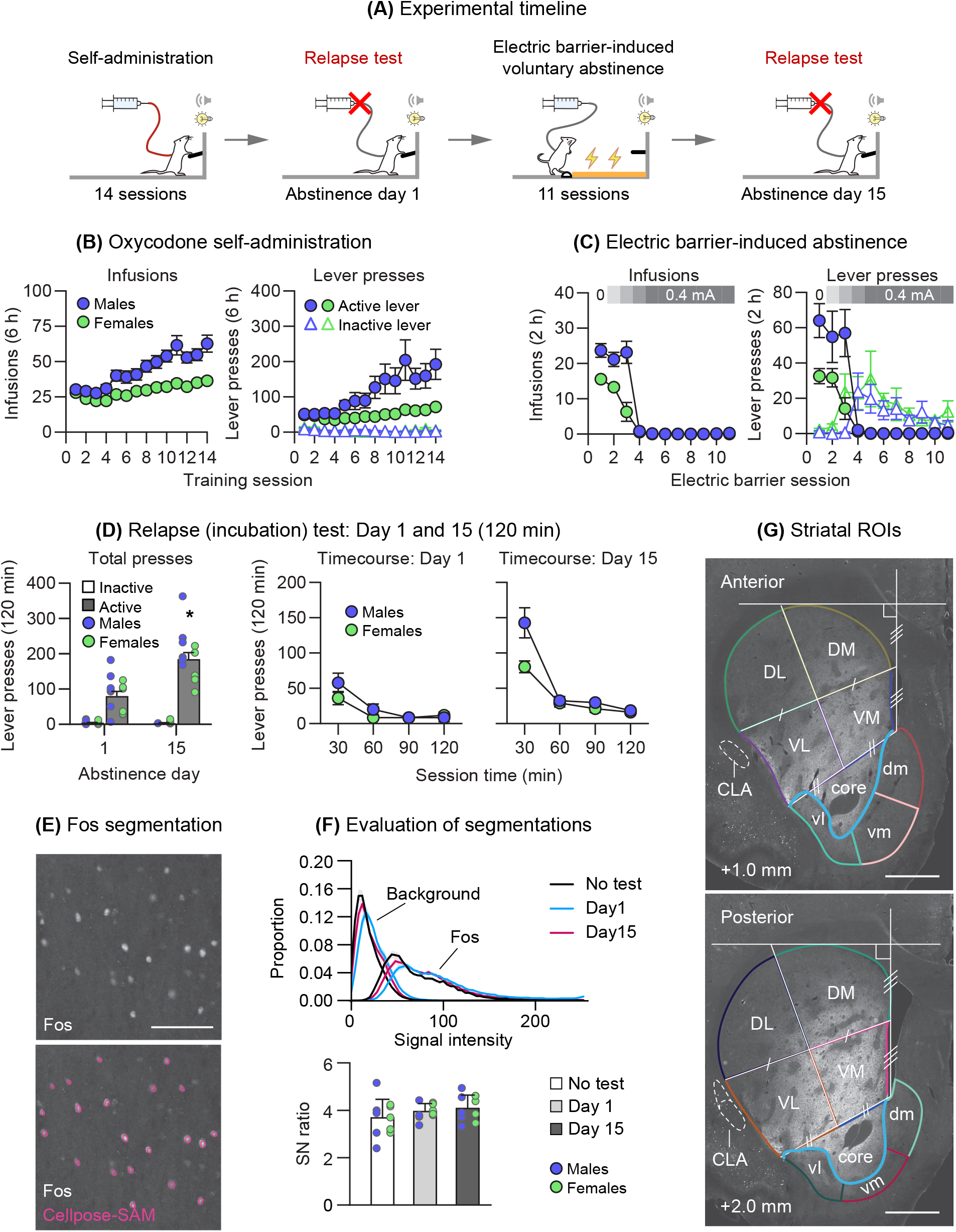
Incubation of oxycodone seeking — Behavioral and histological validation. **(A)** Timeline of behavioral experiment. **(B)** Mean ± SEM number of infusions, active and inactive lever presses during the oxycodone self-administration training for males and females (total n = 28, 14 males, 14 females). **(C)** Mean ± SEM number of infusions, active and inactive lever presses during the electric barrier phase for males and females (total n = 14, 7 males, 7 females). **(D)** Relapse (incubation) tests on abstinence day 1 and day 15 after electric barrier-induced abstinence phase: Left: Mean ± SEM number of active lever presses for males and females during the 120-minute test sessions; Right: Time course of active lever presses for males and females during the 120-minute test sessions, in 30-minute time bins. **(E)** Representative Fos segmentation. Outlines in magenta indicate segmentation outputs. **(F)** *Above*: Histograms of mean signal intensity distributions for background pixels (excluding segmented regions) and per-nucleus mean Fos intensities across No test, Day 1, and Day 15 groups. Upper and lower bounds show SEMs for each group. *Below*: Mean ± SEM signal-to-noise ratios for No test, day 1, and day 15 groups. **(G)** Representative anterior and posterior striatal ROIs based on Nissl, calbindin, and geometric definitions. The blue line indicates the calbindin-based ACB core-shell boundary. * Different from Day 1, p < 0.05. Scale bars: 100 µm **(E)**, 1 mm **(G)**. See **Table 1** for abbreviations.

**Figure 14.**
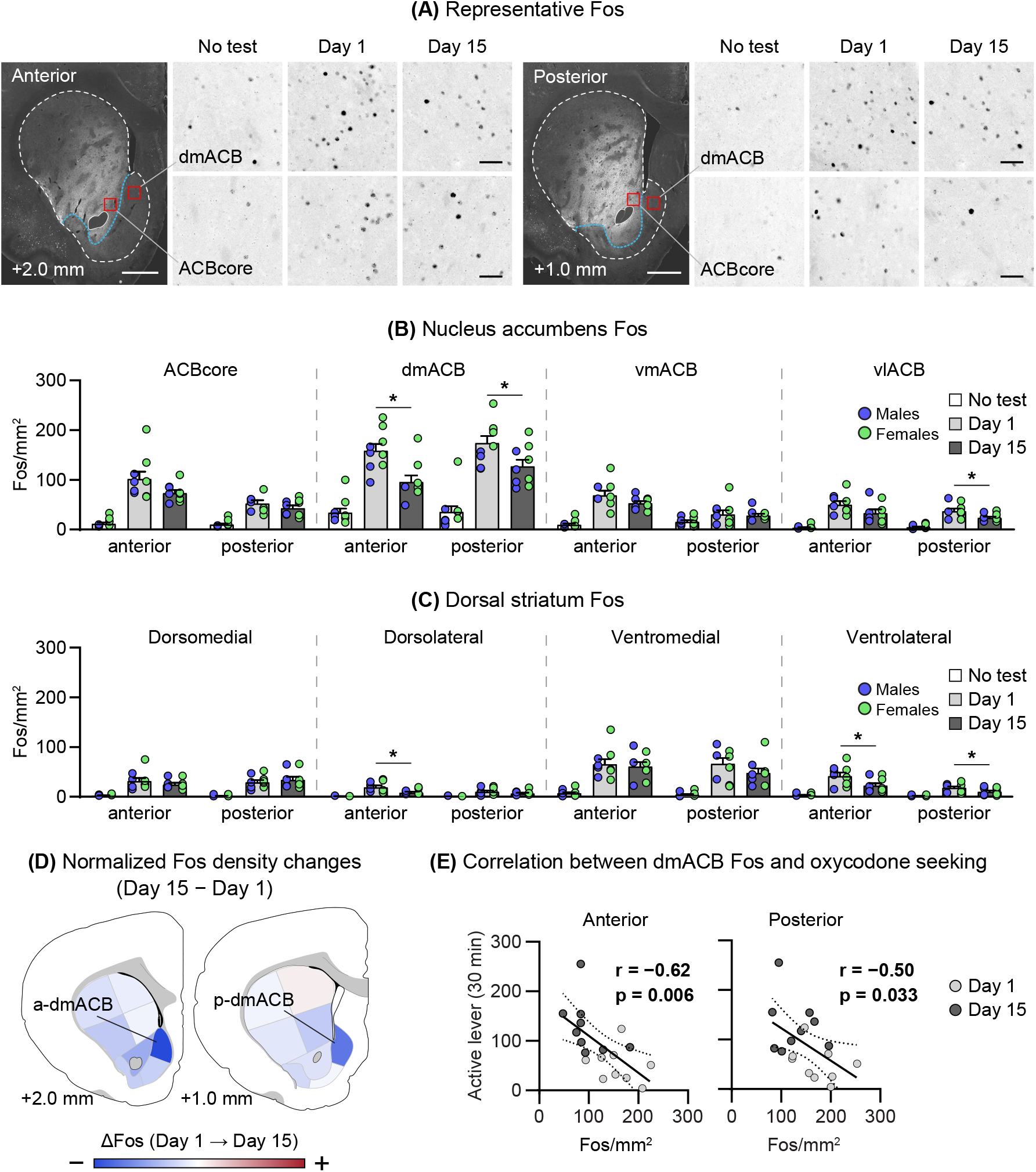
Striatal Fos expression after incubation of oxycodone seeking. **(A)** Representative calbindin image and Fos immunoreactivity (ir) in the dmACB and ACBcore for No test, Day 1 test, and Day 15 test groups. **(B)** Mean ± SEM Fos-ir per mm^2^ in nucleus accumbens regions. **(C)** Mean ± SEM Fos-ir per mm^2^ in dorsal striatum regions. **(D)** ROI representation of Fos density changes between Day 1 and Day 15 test groups. Color scale is normalized to the largest absolute value. **(E)** Scatterplots comparing regional Fos expression with active lever responses from the first 30 min of Day 1 and Day 15 relapse tests. Bold lines show linear regression and dashed lines show upper and lower 95% confidence intervals for combined Day 1 and Day 15 test groups. * Different from Day 1, p<0.05. Scale bars: 50 µm (insets in **A**), 1 mm **(A)**. See **Table 1** for abbreviations. effects and interactions **(Table 2)**.

**Figure 15.**
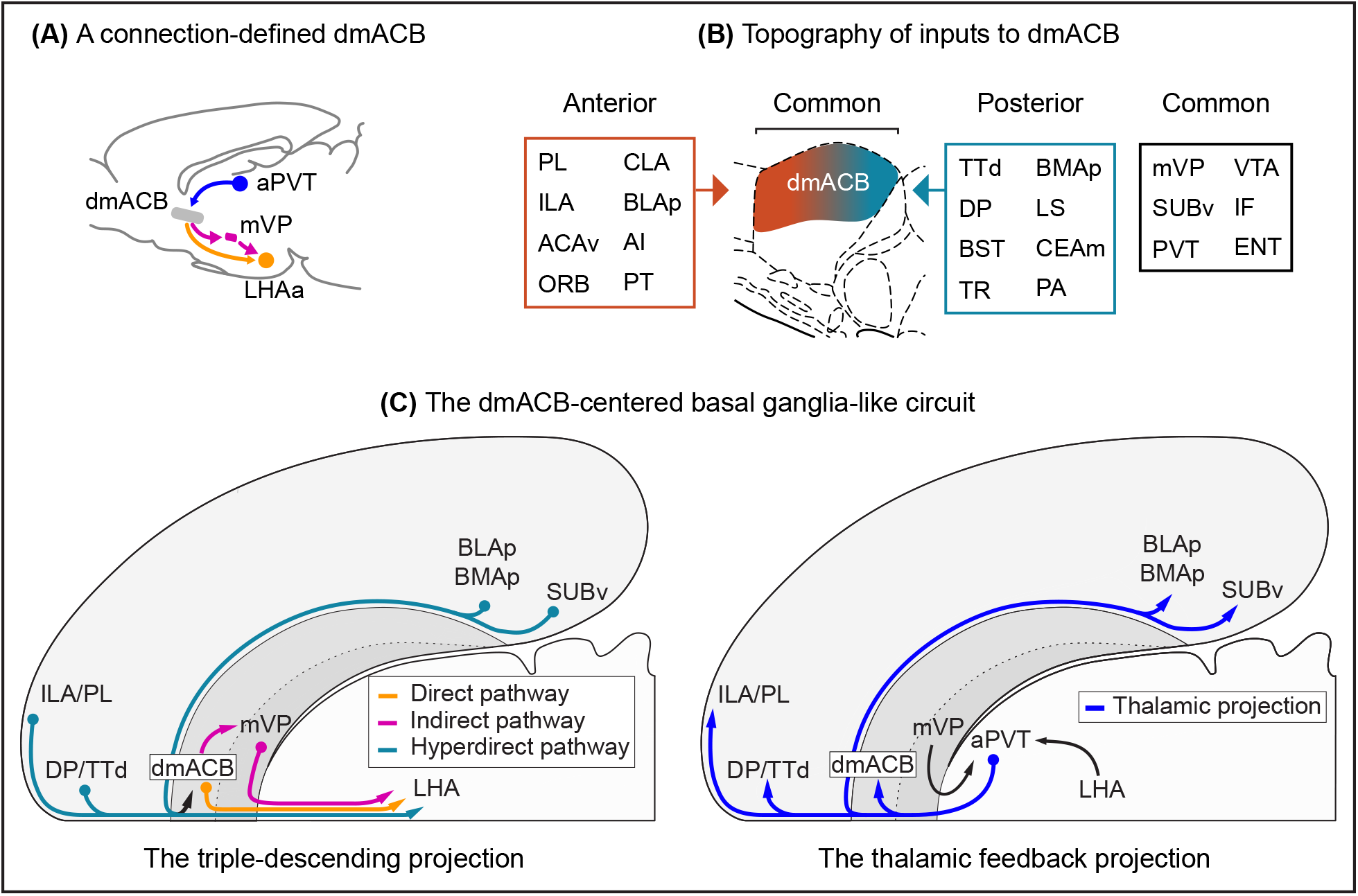
Summary of anatomical findings. **(A)** Diagram of pathways used for connection-based definition of dmACB. **(B)** Summary of main forebrain inputs to anterior and posterior dmACB. **(C)** The updated dmACB circuit follows the general basal ganglia architecture outlined by Swanson (2000) and builds on the dmACB circuit described by Thompson and Swanson (2010). *Left:* A triple-descending projection formed by projections from several cortical areas, dmACB, and mVP that converge on the LHA. *Right:* A thalamic feedback system formed by mVP and LHA inputs to PVT. PVT then projects to dmACB and each cortical node of the circuit (Figs. 9–12; also see Vertes (2008)), completing a distributed closed-loop circuit at the mesoscale. See **Table 1** for abbreviations.

We also found that incubation of oxycodone seeking after electric barrier-induced abstinence was associated with reduced numbers of Fos-activated neurons in the dmACB. Additionally, regardless of the abstinence duration, the number of Fos-positive neurons in the dmACB was inversely associated with active lever pressing during the relapse tests **(Fig. 14)**. This negative correlation was specific to the dmACB and was not observed in other accumbens or dorsal striatum subregions.

Together, these findings indicate that the dmACB is not simply part of a shell gradient, but a distinct anatomical and functional unit within the ventral striatum. We discuss these findings below.

### The dmACB-centered basal ganglia loop

Conceptually, the dmACB-centered circuit aligns with canonical models of basal ganglia organization. Classical frameworks propose parallel, segregated cortico-striato-pallido-thalamocortical loops that support functionally specialized processing (Alexander et al., 1986). The circuit we describe here **(Fig. 15C)** conforms to this architecture and incorporates hypothalamic and thalamostriatal connections, consistent with more recently elaborated models (Barbier et al., 2020; Swanson, 2000).

The dmACB integrates convergent cortical and thalamic inputs and projects to LHAa either directly or indirectly via the mVP (**Fig. 15C**, left panel). The mVP and LHAa, in turn, project to aPVT, which sends thalamostriatal **(Fig. 2)** and thalamocortical **(Figs. 9–12)** outputs, completing a mesoscale closed-loop circuit (Li & Kirouac, 2008, 2012; Thompson & Swanson, 2010) (**Fig. 15C**, right panel). This organization positions the dmACB, at a mesoscale level of analysis, as the striatal node of a basal ganglia loop, an idea that warrants confirmation at the synaptic level. Moreover, our anatomical characterizations and computational analyses suggest that the dmACB represents a discrete, connectionally segregated subdomain within the broader accumbal shell organization (Groenewegen et al., 1993; Thompson & Swanson, 2010; Zahm & Heimer, 1993).

Molecular data further support this interpretation. Recent studies identify a spatially clustered population of “hybrid” neurons within dmACB co-expressing D1 and D2 dopamine receptors, which are segregated from the canonical D1- and D2-receptor expressing populations in manifold space (Gagnon et al., 2017; Gokce et al., 2016; He et al., 2021; Saunders et al., 2018; Stanley et al., 2020). Transcriptomic analyses similarly isolate a Tac3-expressing interneuron cluster in human dmACB (Garma et al., 2024), which appears evolutionarily conserved (Corrigan et al., 2025) and is expressed in primate dmACB (He et al., 2021). Genes associated with these D1-D2 hybrid clusters (e.g., Stard5, Dner, Stra6, Tshz1, and Tac2) are also selectively expressed along the anteroposterior extent of dmACB (Stanley et al., 2020). In mice, D3 receptor-expressing neurons within the dmACB also project to LHA (Enriquez-Traba et al., 2024). Together, these molecular features and our anatomical results support the notion that the dmACB represents a discrete longitudinal striatal module with unique gene-expression and connectivity profiles.

Within the dmACB module, we identified a clear anterior-posterior differentiation. Both the anterior and posterior dmACB receive common inputs from SUBv, mVP, the full extent of PVT, and a ventromedial subdomain of the VTA. These shared inputs provide independent support for the connection-defined organization of the dmACB column and identify candidate upstream components of the dmACB-centered loop. Beyond these common connections, however, the afferent organization diverges. The anterior dmACB receives projections from PL, ILA, ACAv, ORB, BLAp, anterior CLA, AI, and the anterior PT, whereas the posterior dmACB receives projections from TTd, DP, BST, TR, BMAp, LS, PA, and to a lesser extent, from the CEAm **(Fig. 15B)**. Our finding that amygdalar and BST projections preferentially innervate the posterior dmACB is consistent with previous studies in rats (Dong & Swanson, 2003, 2006; Petrovich, 1997; Petrovich et al., 1996) and mice (Hintiryan et al., 2021). Consistent with those studies, we also found that amygdalar and BST projections were almost exclusively ipsilateral, with minimal or no labeling in the contralateral hemisphere (data not shown). In contrast, a recent study in mice reported a prominent interhemispheric amygdalostriatal pathway (Tian et al., 2024). The reasons for this discrepancy are unclear.

Together, our findings indicate that the dmACB column comprises partially segregated anterior and posterior subsystems that integrate distinct corticolimbic information streams while converging onto a common downstream network.

These divergent anterior-posterior inputs to the dmACB may explain previously reported functional differences within this region. Several studies have described opposing behavioral effects depending on whether pharmacological manipulations targeted the anterior or posterior dmACB (Hawks et al., 2025; Peciña & Berridge, 2005; Pirino et al., 2024; Reynolds & Berridge, 2001, 2002). Our data suggest that these differences arise from modulation of distinct subsystems embedded within a common dmACB circuit. For example, pharmacological manipulations that elicit positive valence (“hedonic”) responses in anterior dmACB but aversive responses in posterior dmACB (Castro & Berridge, 2017) align with the distinct cortical and limbic afferent organization identified here.

Recent cell type-specific recordings further support functional differentiation along the anterior-posterior axis of dmACB. In mice, calcium activity in Drd1-expressing neurons decreased following liquid reward consumption in anterior, but not posterior, dmACB (Marinescu et al., 2026). Consistent with this finding, optogenetic stimulation of anterior, but not posterior, dmACB Drd1-expressing neurons reduced reward consumption. In contrast, Drd1-expressing neurons in both anterior and posterior dmACB increased activity following unexpected footshocks and tail suspension, and stimulation of either region produced real-time place aversion (Marinescu et al., 2026). Together, these findings suggest that the anterior and posterior dmACB comprise of partially specialized behavioral subsystems while retaining a shared role in processing aversive or threatening stimuli, potentially through the common dmACB circuit identified here.

We found evidence suggestive of compartmental organization within the dmACB based on PHAL labeling from aPVT and CTB labeling from LHAa. Although striosomes are typically identified by mu-opioid receptor (MOR) or calbindin immunolabeling (Liu & Graybiel, 1992; Morigaki et al., 2020), neither marker clearly delineates compartments in dmACB. Calbindin distinguishes striosomes from matrix in the dorsal striatum but is largely absent from the dmACB **(Fig. 2)**, whereas MOR is expressed in surrounding matrix (Morigaki et al., 2020). Similarly, prepronociceptin, a recently proposed striosome marker, does not clearly distinguish compartments in ACB shell. Candidate markers for dmACB striosomes include Foxp2 and Tshz1 (Takahashi et al., 2003; Xiao et al., 2020). Nevertheless, our findings, and those of others (Contesse et al., 2025), support the existence of compartmental organization within the dmACB and suggest that the circuit described here is organized into compartment-specific subsystems.

Together, our findings suggest that the dmACB is best conceptualized as a longitudinally organized striatal column that is distinct from the surrounding ACB and integrates separate cortical and limbic inputs into a shared hypothalamic-thalamic output pathway. Future work is required to determine how anterior–posterior and compartmental subsystems interact within this shared circuit and how their coordinated or dissociable activity controls motivated behavior.

### dmACB activation and incubation of oxycodone seeking

In our initial characterization of a role for dmACB in motivated behavior, we used Fos immunostaining as a proxy for neuronal activation in anterior and posterior dmACB during incubation of oxycodone seeking following electric barrier-induced abstinence. We also compared the numbers of Fos-labeled cells in dmACB with those within neighboring ventral and dorsal striatal regions.

We found that dmACB activation in both anterior and posterior subregions inversely correlated with relapse responding: The numbers of Fos-labeled cells decreased in association with the relapse test after electric barrier-induced voluntary abstinence. This result appears to contrast with prior studies showing that increased glutamate-mediated activity in both ACB core and shell promotes incubation of cocaine, methamphetamine, and oxycodone seeking after homecage forced abstinence (Chow et al., 2025; Conrad et al., 2008; Dong et al., 2017; Wolf, 2025). However, we should be cautious in generalizing mechanisms of incubation from forced abstinence to incubation after electric barrier-induced abstinence. In our previous work, inactivating the ventral subiculum inhibited incubation after electric barrier-induced abstinence but had no effect on incubation after forced abstinence, suggesting that partially dissociable circuits mediate incubation after these two forms of abstinence (Fredriksson et al., 2023; Negishi et al., 2025). Additionally, under our experimental conditions, increased numbers of Fos-labeled cells during the relapse test likely reflects multiple processes in addition to promoting or inhibiting drug seeking, including operant responding, extinction learning, reward omission, and drug context exposure. Moreover, because we detected Fos with an antibody selective for its Ser^32^-phosphorylated form, both the lower numbers of Fos-labeled dmACB cells after the Day 15 test and their negative correlation with oxycodone seeking may reflect concurrent changes in Fos induction and phosphorylation state. Finally, we did not conduct experiments to identify striatal cell types that expressed Fos. Therefore, without cell type attribution and causal manipulation of the relapse-associated Fos-expressing neurons (Cruz et al., 2013), the observed negative correlation between dmACB activity and incubation of oxycodone seeking should be interpreted with caution.

Our correlational finding that dmACB activation inversely correlates with oxycodone seeking suggests that neuronal activity in this region inhibits relapse-related behavior, which is an unexpected possible outcome given the broader literature implicating ACB shell activity in drug reward and motivated behavior (Everitt & Robbins, 2005; Ikemoto, 2007; Wise, 2004). However, our results are consistent with several studies indicating that neuronal activation in ACB shell (including the dmACB region we characterized here) inhibits appetitive behaviors, including feeding (Kelley, 2004), reinstatement of drug seeking (Gibson et al., 2019), and other reward-related behaviors (Nicola, 2007).

In these studies, microinjecting pharmacological inhibitors of ACB shell neuronal activity (e.g., with GABAA receptor agonists or glutamate receptor antagonists) increased feeding and drug seeking, whereas pharmacological stimulators of ACB shell activity produced opposite effects (Di Ciano et al., 2008; Kandov et al., 2006; Kelley, 2004; Maldonado-Irizarry et al., 1995; Millan & McNally, 2011; Peters et al., 2009; Peters et al., 2008; Stratford & Kelley, 1999; Urstadt et al., 2013). Further supporting this notion, ACB shell microinjections of dopamine D2 receptor antagonists, which increase striatal Fos expression after systemic injections (Dragunow et al., 1990; Robertson et al., 1991), reduce cocaine priming-induced reinstatement of drug seeking (Anderson et al., 2006; Schmidt et al., 2005).

Recent optogenetic and cell-type-specific studies further support these findings. Dopamine D1 receptor-expressing neurons in ACB shell, including dmACB, are critical for inhibition of alcohol seeking (Gibson et al., 2018) and social- and feeding-related behaviors (Contesse et al., 2025) via projections to LHA. Additionally, Pomrenze et al. (2024) showed that dmACB *Tshz1*-expressing cells are activated during naloxone-precipitated opioid withdrawal, and chemogenetic inhibition of these cells decreases opioid withdrawal-induced conditioned place aversion.

Together, classical reversible inactivation/activation studies and modern cell-type and projection-specific manipulations indicate that dmACB activity constrains appetitive-motivated behaviors and promotes aversive-motivated behaviors. Our data showing an inverse correlation between the numbers of Fos-labeled cells in dmACB and relapse to opioid seeking in a rat model supports this conclusion.

## Conclusion

Our anatomical results indicate that the dmACB is a unique ACB subregion, located primarily within the classical ACB shell (Groenewegen et al., 1999). We showed that the dmACB forms a longitudinal structure extending along the anterior-posterior axis, defined by convergent anatomical connections with mVP, LHAa, and aPVT. Although the dmACB is defined by a common set of connections, it receives divergent cortical and subcortical inputs along the anterior-posterior axis. Using a new model for incubation of oxycodone-seeking electric barrier-induced abstinence, we also showed that dmACB activation was unexpectedly inversely associated with this form of incubation. We propose that dmACB is a node in a discrete basal ganglia loop whose activity constrains opioid relapse following abstinence induced by the adverse consequences of drug seeking.

## Data and materials availability

All BM4 atlas maps were modified and/or reproduced with permission under the conditions outlined by a Creative Commons BY-NC 4.0 license (https://creativecommons.org/licenses/by-nc/4.0/). The behavioral and neuroanatomical data generated during this study are openly available in the Harvard Dataverse repository at https://doi.org/10.7910/DVN/DZQTQD. The Python code and associated data-processing procedures are described in the Methods; the scripts are available the scripts are available at https://github.com/ken-negishi/dmACB_project.

## Acknowledgments

This research was supported [in part] by the Intramural Research Program of the National Institutes of Health (NIH). The contributions of the NIH author(s) are considered Works of the United States Government. The findings and conclusions presented in this paper are those of the author(s) and do not necessarily reflect the views of the NIH or the U.S. Department of Health and Human Services. We thank Alexander Friedman for helpful comments. We also thank Alexandro Arnal for early discussions and preliminary code that informed the computational strategy for axon segmentation and kernel density estimation used in this study.

## Author Contributions

All authors contributed to different aspects of the study, including the design and performance of the research, data analysis, and write-up of the paper.

## Ethics approval statement

Procedures were approved by local Ethical Committee of the Intramural Research Program of NIDA, NIH, and the Institutional Animal Care and Use Committee of The University of Texas at El Paso.

